# Host Vesicle Fusion Proteins VAPB, Rab11b and Rab18 Contribute to HSV-1 Infectivity by Facilitating Egress through the Nuclear Membrane

**DOI:** 10.1101/088633

**Authors:** Natalia Saiz-Ros, Rafal Czapiewski, Andrew Stevenson, Ilaria Epifano, Selene K. Swanson, Marion McElwee, Swetha Vijayakrishnan, Christine A. Richardson, Charles Dixon, Lior Pytowski, Martin W. Goldberg, Laurence Florens, Sheila V. Graham, Eric C. Schirmer

**Author notes:** Correspondence to: Eric Schirmer, Wellcome Trust Centre for Cell Biology, University of Edinburgh, Kings Buildings, Michael Swann Bldg., room 5.22, Max Born Crescent, Edinburgh EH9 3BF UK.

## Abstract

The herpesvirus process of primary envelopment and de-envelopment as viral particles exit the nucleus has been for many years one of the least understood steps in the virus life cycle. Though viral proteins such as pUL31, pUL34, pUS3 and others are clearly important, these are likely insufficient for efficient fusion with the nuclear membrane. We postulated that host nuclear membrane proteins involved in virus nuclear egress would move from the inner to outer nuclear membranes due to membrane fusion events in primary envelopment and de-envelopment and then diffuse into the endoplasmic reticulum. Membrane fractions were prepared enriched in the nuclear envelope or the endoplasmic reticulum with and without HSV-1 infection and analyzed by mass spectrometry, revealing several vesicle fusion proteins as candidates in the viral nuclear egress pathway. Knockdown of three of these, VAPB, Rab11b, and Rab18, significantly reduced titers of released virus while yielding nuclear accumulation of encapsidated particles. Antibody staining revealed that VAPB visually accumulates in the inner nuclear membrane during HSV-1 infection. VAPB also co-localizes at early time points with the viral pUL34 protein known to be involved in nuclear egress. Most strikingly, VAPB was also observed on HSV-1 virus particles by immunogold labelling electron microscopy. Thus, these data reveal several new host cell vesicle fusion proteins involved in viral nuclear egress.

**Author Summary:** Human herpesviruses are associated with common human diseases such as chicken pox, shingles and mononucleosis and infect a wide range of animals making them economically important pathogens for livestock. Herpes simplex virus 1 (HSV-1) is most commonly associated with cold sores, but is also the leading cause of blindness by infection in the Western world. All herpesviruses share many aspects of infection. As large nuclear replicating dsDNA viruses with capsid sizes too large to use the nuclear pores to exit the nucleus, they have evolved a complex mechanism for envelopment and de-envelopment of primary herpesvirus particles, but this critical step in the virus lifecycle remains poorly understood. We have identified several host cell vesicle fusion proteins, VAPB, Rab11b and Rab18 that appear to contribute to this step in the HSV-1 life cycle. VAPB accumulates at the nuclear envelope with the HSV-1 pUL34 protein important for viral nuclear egress. Knockdown of any of these vesicle fusion proteins reduces viral titers, further arguing that they are important for nuclear egress. As there appears to be a specific subset of vesicle fusion proteins involved in viral egress, they could possibly represent novel targets for therapeutic interventions.

## Introduction

Herpesviruses are large, enveloped DNA viruses that replicate their genomes and package the nascent progeny viral genomes into capsids in the nucleus while final maturation occur in the cytoplasm. Herpesvirus capsids are too large (~120 nm) to pass through the host cell nuclear pore complexes [1]. Therefore to escape the nucleus they must go through both the inner (INM) and outer (ONM) membranes of the double membrane nuclear envelope (NE) [2]. Work by several labs over many years has captured images of virus particles in various stages of nuclear egress, suggesting a process where tegumented capsids enter the NE lumen after obtaining a primary envelope from the INM and then these particles subsequently exit the lumen by fusing with the ONM and leaving the primary envelope behind [3-6].

Several viral proteins have been identified that are important for this process. First, before capsids can access the nuclear membrane the nuclear lamin meshwork must be dissolved. While many details of the mechanism remain to be worked out, it is clear that HSV-1 pUL31 and pUL34 proteins can recruit host kinases to locally dissolve the nuclear lamina allowing access of nucleocapsids to the INM [7-11]. pUL31 and pUL34 may have additional roles as their overexpression can promote vesiculation both *in vitro* and *in vivo* [12, 13]. Though its specific function is not clear, pUS3 clearly contributes to nuclear egress as its deletion results in accumulation of primary enveloped particles in the NE lumen [14-16]. Additionally, though also part of the final (secondary) envelope, herpesvirus glycoproteins gB and gH contribute to nuclear egress as their combined knockdown results also in accumulation of primary enveloped particles in the NE lumen [17, 18]. Nonetheless, the study of vesicle fusion processes in cells has revealed them to be highly complex and to involve a much larger number of players [19, 20] and herpesviruses are well known for co-opting host proteins [21-23] and so we postulated that nuclear membrane proteins might contribute to this process.

Nuclear membrane proteins enter the nucleus after synthesis in the endoplasmic reticulum (ER) by lateral diffusion through the peripheral channels of the NPC [24, 25]. These are narrow ~10 nm channels that would allow passage for up to a ~60 kDa globular protein. According to the lateral diffusion-retention hypothesis, once inside the nucleus interactions with the nuclear lamina largely immobilize transmembrane proteins in the INM [26]. However, when in the ONM such proteins can rapidly diffuse throughout the endoplasmic reticulum [27-29]. Thus, it might be expected that upon fusion with the ONM during de-envelopment, the host proteins facilitating this process and primary envelope proteins released would diffuse into the ER.

With this idea in mind we prepared membrane samples enriched in NEs or the ER (microsomes) from mock or HSV-1 infected cells and searched for proteins that changed abundance in the ER with infection. Intriguingly, we found several proteins involved in vesicle fusion exhibiting such changes. Knockdown of three of these, VAPB, Rab11b and Rab18 yielded reduced titers of released virus and accumulation of virus particles in the nucleus. VAPB moreover accumulated at the NE early in infection, could be observed to co-localize with the virus pUL34 protein, and most strikingly was observed by immunogold labelling electron microscopy on nucleus associated virus particles. Together these data clearly show that VAPB, Rab11b and Rab18 are important for nuclear egress.

## Materials and Methods

### Cells and viruses

HeLa and U20S cells were grown in Dulbecco’s modified Eagles medium (DMEM, Invitrogen) with 10% fetal calf serum, 50 ¼g/ml penicillin and 50 ¼g/ml streptomycin at 37°C in 5% CO_2_. The HSV-1 wild type strain 17+ was used in all experiments unless specifically noted. For certain experiments an HSV-1 strain expressing RFP fused to VP26 (kindly provided by Frazer Rixon, MRC Virology Unit, University of Glasgow, Glasgow, UK) and the HSV-1 strain vBSGFP27 expressing GFP fused to WT ICP27 under its own promoter [30] was used. HSV-1 strains were routinely propagated and titrated in U2OS cells. Cells were infected at indicated times allowing virus to be adsorbed onto cells for one hour and incubated at 37°C. Viruses were used at multiplicity of infections (MOIs) given in the figure legends.

### Preparation of fractions

Nuclear envelopes (NEs) and microsomal membranes (MMs) were isolated using well-established procedures [31, 32]. In brief, nuclei were first isolated from HeLa cells by hypotonic lysis in 10 mM HEPES pH 7.4, 1.5 mM MgCl_2_, 10 mM KCl, 2 mM DTT using a ‘tight’ dounce homogenizer (Wheaton). Nuclei were pelleted at 1000xg for 10 min to separate them from small vesicles and mitochondria that require higher speeds to pellet. To float/remove contaminating membranes, nuclei were resuspended in SHKM (50 mM HEPES pH 7.4, 25 mM KCl, 5 mM MgCl_2_, 1 mM DTT and 1.8 M sucrose) and pelleted through a 5 ml 2.1 M sucrose cushion in a SW28 swinging bucket rotor (Beckman) at 4°C for 2 h at 82,000 x g. NEs were then prepared from isolated nuclei by two rounds of digestion with DNase and RNase in 0.3 M sucrose, 10 mM HEPES pH7.4, 2 mM MgCl_2_, 0.5 mM CaCl_2_, 2 mM DTT for 20 min followed by layering onto the same buffer with 0.9 M sucrose and centrifugation at 6,000 x g for 10 min at 4°C.

Microsomes were isolated following similar established procedures. In brief, after removing nuclei as for NE preparations, 0.5 mM EDTA was added to inhibit metalloproteinases and mitochondria and other debris from post-nuclear supernatants were also removed by pelleting at 10,000 x g for 15 min. The supernatant was made to 2 M sucrose with SHKM and then overlaid with 1.86 M and 0.25 M sucrose layers. This was then subjected to centrifugation in a SW28 swinging bucket rotor (Beckman) at 4°C for 4 h at 57,000 x g to float microsomes. The material between the 1.86 and 0.25 M layers was then diluted 4-fold with 0.25 M SHKM and pelleted at 152,000 x g in a type 45 Ti rotor (Beckman) for 1 h.

NEs were extracted with 0.1 N NaOH, 10 mM DTT and pelleted by centrifugation at 150,000 x g for 30 min and washed 3x in sterile H_2_O. MMs samples were washed in sterile H_2_O without NaOH extraction. Samples were used for mass spectrometry analysis. HSV-1 infected MMs pellets were prepared and used for EM imaging.

### Mass spectrometry

Membrane pellets were resuspended in 30 ¼l of 100 mM Tris-HCl pH 8.5, 8 M Urea, then brought to 5 mM Tris(2-Carboxylethyl)-Phosphine Hydrochloride (TCEP) and incubated 30 min at room temperature. Next this was reduced and alkylated with 10 mM chloroacetamide (CAM; Sigma) and incubated in the dark for 30 min followed by addition of Endoproteinase Lys-C (Roche) at 0.1 mg/ml and incubated 6 h at 37°C. The solution was diluted for 2 M Urea with 100 mM Tris-HCL pH 8.5 and 2 mM CaCl_2_ and 0.1 mg/ml Trypsin added and incubated at 37°C overnight. Formic acid was then added to 5% to quench reactions and samples were centrifuged to remove large undigested material [33].

Samples were then individually analyzed by Multidimensional Protein Identification Technology (MudPIT) [34, 35]}. Peptide mixtures were pressure-loaded onto 250 μm fused silica microcapillary columns packed first with 3 cm of 5-μm Strong Cation Exchange material (Luna; Phenomenex), followed by 1 cm of 5-μm C18 reverse phase (Aqua; Phenomenex). Loaded 250 μm columns were connected using a filtered union (UpChurch) to 100 μm fused-silica columns pulled to a 5 μm tip using a P 2000 CO_2_ laser puller (Sutter Instruments) and packed with 9 cm of reverse phase material. The loaded microcapillary columns were placed in-line with a Quaternary Agilent 1100 series HPLC using a 10-step chromatography run over 20 h using a flow rate of 200–300 nl/min. The application of a 2.5 kV distal voltage electrosprayed the eluting peptides directly into a LTQ linear ion trap mass spectrometer (Thermo Scientific) equipped with a custom-made nano-LC electrospray ionization source and full MS spectra were recorded on the peptides over a 400 to 1,600 m/z range, followed by five tandem mass (MS/MS) events sequentially generated in a data-dependent manner on the first to fifth most intense ions selected from the full MS spectrum (at 35% collision energy). Dynamic exclusion was enabled for 120sec.

RAW files were extracted into ms2 file format [36] using RawDistiller v. 1.0 [37]. MS/MS spectra were queried for peptide sequence information using SEQUEST v.27 (rev.9) [38] against 55,691 human proteins (non-redundant NCBI 2014-02-04 release), plus 162 sequences from usual contaminants (e.g. human keratins…) and 77 NCBI RefSeq HSV1 proteins. To estimate false discovery rates, each non-redundant protein entry was randomized and added to the database bringing the total search space to 111,524 sequences. MS/MS spectra were searched without specifying differential modifications, but +57 Da were added statically to cysteine residues to account for carboxamidomethylation. No enzyme specificity was imposed during searches, setting a mass tolerance of 3 amu for precursor ions and of ±0.5 amu for fragment ions.

Results from different runs were compared using DTASelect and CONTRAST [39] with criterion of DeltCn ≥ 0.08, XCorr ≥ 1.8 for singly-, 2.5 for doubly-, and 3.5 for triply-charged spectra, and a maximum Sp rank of 10. Peptide hits from all analyses were merged to establish a master list of proteins (Tables S1). Identifications mapping to shuffled peptides were used to estimate false discovery rates. Under these criteria the final FDRs at the protein and peptide levels were on average 0.72% and 0.24%, respectively.

To estimate relative protein levels, distributed normalized spectral abundance factors (dNSAFs) were calculated for each non-redundant protein (Table S1), as described in [40]. The complete MudPIT mass spectrometry dataset (raw files, peak files, search files, as well as DTASelect result files) can be obtained from the MassIVE database via <u>ftp://MSV000079886@massive.ucsd.edu</u> with a password NSR7095. The ProteomeXchange accession number for this dataset is PXD004519.

### Bioinformatics analysis

To detect proteins enriched in HSV-1 infected MMs, the ratio of HSV-1 infected MMs versus mock-infected MMs was calculated according to relative abundance determined by dNSAF values. To choose only proteins present in the NE before infection, the ratio of mock-infected NE versus mock-infected MMs was similarly calculated. Only proteins with a HSV-1 infected MMs: mock-infected MMs ratio higher than 1.3 and detected in the mock infected NE were selected.

Gene ontology (GO) analysis was performed in Ensembl BioMart and biologically interesting categories were represented either as a piechart or shown as bar plots in %. Biologically interesting GO-terms and their corresponding child terms were retrieved from the mySQL database <u>http://amigo.geneontology.org</u>[41]. We used the following GO terms: “vesicle-mediated transport” (GO:0016192) “nucleocytoplasmic transport” (GO:0006913), “membrane organization” (GO:0061024), “regulation of protein phosphorylation” (GO:0001932). To plot MS data pie charts, the relative proportions of the various classes were calculated from the relative abundances of the genes identified from each GO category, and this was compared to a pie chart compiled from all the genes in the human genome considering the same categories.

### siRNA knockdown of vesicle fusion proteins

HeLa cells were plated in 12 well plates at 5.5x10^4^ cells per well and were transfected with Rab1a, Rab11b, VAPB, Rab18 and Rab24 with 50 nM total small interfering RNA (siRNA) using JetPrime Polyplus Transfection reagent according to the manufacturer’s instructions. The Rab1a, Rab11b, Rab18 and Rab24 siRNAs were all SMARTpools from Dharmacon. VAPB single siRNA oligos were obtained from SIGMA and have been previously described [42]. The control siRNA was a scrambled sequence (5’-CGUACGCGGAAUACUUCGA-3’) of an siRNA to firefly luciferase that has frequently been used as a generic control in studies where multiple proteins are targeted. At 48 h after the final transfection, the total final protein was extracted for Western blot analysis or cells were infected with HSV-1 WT strain 17+ at MOI 10 to determine viral replication. For these, samples were harvested at 16 hpi and supernatant collected for plaque assays.

### Cloning of VAPB, RAB11b and Rab1A

Primers for PCR amplifying VAPB, RAB11B and RAB1A were designed for in Fusion cloning (Clontech) into pCDNA3.1 using the following software: (<u>http://www.clontech.com/US/Products/Cloning_and_Competent_Cells/Cloning_Resources/Online_In-Fusion_Tools</u>).The primers were designed to retain the *Eco* RI and *Bam* HI sites in the multiple cloning site of pCDNA3.1. For initial cloning of VAPB, RAB11B and RAB1A PCR was carried out using the appropriate primer set (Eurofins) and CloneAmp HiFi PCR Premix (Clontech). The template was cDNA that had been synthesized (Maxima kit with dsDNAse ‐Thermofisher) from RNA that had been extracted from differentiated NIKS16 cells. PCR products were treated with cloning enhancer (Clontech) and mixed with pCDNA3.1 (which had been previously digested with *Eco*RI and *Bam*HI (NEB) to linearize the vector) and cloned using the In-Fusion HD Cloning Kit (Clontech). Reactions were transformed into *E. coli* Stellar Competent Cells and selected on 100 ¼g/ml Ampicillin (Sigma). Plasmid DNA was prepared (Qiagen QIAprep Spin Miniprep Kit) and plasmids digested *Eco*RI and *Bam*HI to confirm insert. Clones were then verified by DNA sequencing (Eurofins).

### Mutagenesis of VAPB

For site directed mutagenesis of VAPB ((DNA sequence CTTGGCTCTGGTGGTT change to GTTTGCACTTGTCGTG) an inverse PCR was carried out on pCDNA3.1VAPB using primer set VAPB SDMfw1 and VAPBSDMrv1 with CloneAmp HiFi PCR Premix (Clontech). The PCR product was treated with cloning enhancer (Clontech) and the In-Fusion HD Cloning Kit (Clontech). Reactions were transformed into *E. coli* Stellar Competent Cells and selected on 100 ¼g/ml Ampicillin (Sigma). Plasmid DNA was prepared (Qiagen QIAprep Spin Mindiprep Kit) and the clone and mutation were verified by DNA sequencing (Eurofins). Note that mutagenesis was not required for Rab1A and Rab11b rescue constructs because their siRNAs targeted the 3’ untranslated region.

### Rescue experiment

Hela cells expressing the rescue constructs were plated in 12 well plates and transfected the next day with 50 nM of Rab1A, VAPB or Rab11b siRNA; 0.5 ¼g of pCDNA3.1 and 0.5 ¼g pCDNA3.1-GFP plasmid to check the efficiency of the transfection. After 48 h, cells were infected with HSV-1 WT at MOI 10. After 16 hpi, supernatants were collected for plaque assays.

### Viral plaque assay

Cellular culture supernatant was harvested after 16 hpi. Samples were frozen at -80°C, thawed, and sonicated. Tenfold serial dilutions were made from these viral supernatant, and the samples were incubated on confluent U2OS cells for 1h at 37°C. The virus then was aspirated, cells were wash in PBS and overlaid with a 1:1 mixture of 1% (wt/vol) avicel (RC-591 NF) in sterile water–FBS-DMEM. The overlaid samples were cultured for 3 days before the avicel/medium overlay was removed. The cells then were fixed with methanol and stained with crystal violet solution (2% [wt/vol] crystal violet in 40% [vol/vol] methanol) in order to quantify viral plaque formation.

### Fluorescence in situ hybridization

For FISH experiments HeLa cells were cultured on coverslips that were washed in PBS prior to fixation in 4% para-formaldehyde in 1X PBS for 10 min at room temperature. Cells were permeabilized for 6 min with 0.2% Triton-X-100 in PBS, followed by 3 washes in PBS. Cells were next pre-equilibrated in 2X SCC and treated with RNase A (100¼g/ml) at 37ºC for 1 h. Following washing in 2X SCC, cells were dehydrated with a 70%, 85% and 100% ethanol series. Coverslips were then air dried, heated to 70ºC and submerged into 85ºC preheated 70% formamide, 2X SSC (pH 7.0) for 18 min. A second ethanol dehydration series was then performed using -20ºC 70% ethanol for the first step. Coverslips were air dried and 150-300 ng biotin-labelled probe was added in hybridization buffer (50% formamide, 2X SSC, 1% Tween20, 10% Dextran Sulphate) containing 6 ¼g human Cot1 DNA (Invitrogen) and sheared salmon sperm DNA and incubated at 37°C for 24 h in a humidified chamber. Probes were generated from a plasmid encoding HSV-1 gene ICP27 by end labelling. After incubation, the coverslips were washed four times for 5 min each in 4X SSC at 50ºC followed by four times for 5 min each in 0.1X SSC at 65ºC. Coverslips were then pre-equilibrated in 4X SSC, 0.1% Tween-20 and blocked with 4% BSA before incubating for 30 min at room temperature with Alexa Fluor®conjugated-Steptavidin antibodies and 4,6-diamidino-2 phenylindole, dihydrochloride (DAPI) at 2 ¼g/ml. Coverslips were subsequently washed 3 times in 4X SSC, 0.1% Tween-20 at 37ºC and mounted on slides in Vectashield (Vector Labs).

Images from the midplane of the nucleus were acquired on a Nikon TE-2000 microscope using a 1.45 NA 100x objective, Sedat quad filter set, and a CoolSnapHQ High Speed Monochrome CCD camera (Photometrics) run by Metamorph image acquisition software. Images were analyzed using ImageJ, using the DAPI stained image as a mask to determine nuclear area and the total fluorescence intensity in the nucleus and in the whole cell were determined for over 100 cells for each condition. The total cellular fluorescence divided by the nuclear fluorescence is plotted in Figure 6.

### Antibodies and Western blotting

Proteins were fractionated by SDS-PAGE and transferred to nitrocellulose membranes. Membranes were blocked with 5% milk powder in PBS containing 0.1% Triton-X-100 (PBST) for 1 h at RT. Membranes were incubated with primary antibodies (Abs) in the same buffer for 1–2 h at room temperature or overnight at 4°C, washed, and incubated with anti-mouse IgG or anti-rabbit IgG conjugated to horseradish peroxidase for 1 h with shaking. Following washing in PBST, proteins on membranes were visualized either using ECL (GE Healthcare) and exposed to Kodak X-Omat S film or analyzed directly on a LICOR Odyssey imager (LI-COR Biosciences) using antibodies conjugated to fluorescent markers. Abs used were against GM-130 (EP892Y) from Abcam, GAPDH (E1C604-1; 1:500) from Enogene, Calnexin (Stressgen, SPA-860), lamin A and C (3262, 3931 [43]), VAPB (rabbit antibody used in Western blots) was kindly provided by Dr. Christopher C.J. Miller, Rab11b (*NBP2*-15085), Rab18 (IHC-plus ^TM^ LS-B9430), Rab1a (Santa Cruz Biotechnology), Rab24 (BD bioscience), gC (ab6509) and US3 (was kindly provided by Dr. Thomas Mettenleiter and Dr. Barbara Klupp)

### Fluorescence Microscopy

HeLa cells were grown in monolayers on each 13 × 13 mm coverslip overnight, then infected with virus or mock infected for the times stated. Coverslips were washed three times with PBS and fixed with 3.7% formaldehyde for 10 min, then permeabilized with 0.2% Triton X-100 for 5 min at RT for fixation and finally washed three times with PBS. Coverslips were blocked with PBS containing 1% human serum, 1% calf serum and 0.1% BSA for 1 h at room temperature, then incubated with primary Abs diluted in same blocking buffer for 1 h at room temperature: VAPB (mouse monoclonal, Proteintech) and UL34 (chicken antibody kindly provided by Dr. Richard Roller). Coverslips were washed in PBS six times before incubation for 1 h with secondary Abs (Jackson Laboratories, donkey minimal cross-reactivity) diluted in blocking buffer. After washing in PBS six times, cells were incubated with 4’,6-diamidino-2-phenylindole (DAPI; 1:2000) as a nuclear stain and washed with PBS and coverslips were mounted with vectashield mounting medium.

Most images were obtained using a Nikon TE-2000 microscope equipped with a 1.45 NA 100x objective, Sedat quad filter set, PIFOC Z-axis focus drive (Physik Instruments), and CoolSnapHQ High Speed Monochrome CCD camera (Photometrics) run by Metamorph image acquisition software. Image stacks (0.2 μm steps) were deconvolved using AutoquantX. Micrographs were saved from source programs as 12-bit ˙tif files and analyzed with Image Pro Plus software and/ or prepared for figures using Photoshop 8.0.

For super resolution microscopy, cells were prepared and stained similarly except that images were taken using a Zeiss 880 confocal microscope with Airyscan and prepared for figures using Photoshop 8.0.

### Electron Microscopy

MM pellets were washed in sterile H_2_O and fixed with 2.5% glutaraldehyde and 1% osmium tetroxide, then dehydrated through a graded alcohol series and embedded in Epon 812 resin. Thin sections were cut and examined in a JEOL 1200 EX II electron microscope.

To test the localization of viral particles in cells with knockdowns of vesicle fusion proteins, monolayer cultures of HeLa cells were seeded in a 60 mm dish and transfected with siRNAs against VFPs as above. 48 hours later cells were infected with HSV for 16 hpi at an MOI of 10, and subsequently fixed with 2.5% glutaraldehyde in 2% sucrose, 0.05M Cacodylate buffer, pH 7.2 overnight at 4°C. Cells were thoroughly washed with PBS, scraped and pelleted by centrifugation followed by fixation with 1% osmium tetroxide (TAAB Labs, UK) and staining with 2% aqueous uranyl acetate for 1 h at room temperature. Cells were then harvested into PBS and pelleted through 3% low gelling temp agarose (Geneflow) at 45°C. The agarose was set at 4°C and cell pellets were cut into ~1 mm cubes, which were dehydrated through a graded alcohol series (30-100%) and embedded in Epon 812 resin (TAAB Labs, UK) followed by polymerization for 3 days at 65°C. Thin sections of 120 nm were cut with a UC6 ultramicrotome (Leica Microsystems, Germany) and examined with a JEOL 1200 EX II electron microscope and images were recorded on a Gatan Orius CCD camera.

Immunoelectron microscopy was performed using the Tokuyasu (1973) method [44]. Cells were fixed *in situ* with 2% paraformaldehyde, 0.2% glutaraldehyde in PHEM buffer (60 mM Pipes, 25 mM Hepes, 2 mM MgCl_2_, 10 mM EGTA, pH 6.9) for 1 h, washed with PHEM buffer, then scraped off the culture dish. They were pelleted at 200 X g for 2 min and resuspended 0.1% glycine in PBS, pelleted at 400 x *g* for 2 min, resuspended 0.1% glycine in PBS for 15 min, pelleted at 400 x *g* for 2 min, resuspended in 1% gelatin (Dr Oetker) at 37°C for 10 min, pelleted at 400 x *g* for 2 min, resuspended in 10% gelatin for 10 min at 37°C, then replaced on ice. Gelatin embedded pellets were cut into ~5 mm cubes and immersed in 15% PVP (polyvinylpyrrolidone 10K molecular weight, Sigma), 0.17 M sucrose in PBS overnight. Samples were mounted and frozen in liquid nitrogen then sectioned on a cryo-ultramicrotome (UC6 with FC6 cryo-attachment; Leica). Cryosections were thawed, and the gelatin melted at 37°C, washed in 0.1% bovine serum albumin (BSA, Sigma) in PBS, then 1% BSA in PBS for 3 min, followed by overnight incubation with undiluted primary antibody, washed in PBS, incubated with secondary anti– mouse antibody conjugated to 10 nm colloidal gold (BBI solutions). Grids were then washed in PBS, transferred to 1% glutaraldehyde in PBS (5 min), washed in water, and embedded in 2% methyl cellulose containing 0.4% uranyl acetate (Agar Scientific). A Hitachi H7600 TEM was used at 100 kV.

### qPCR

Cells transfected with siRNA against Rab1a, Rab24, Rab11b, VAPB and Rab18 as described above were infected with HSV-1 WT at an MOI of 10. Supernatants or pellets were collected at 16 hpi. Viral DNA was purified from scraped pellet using the DNeasy Blood&Tissue Kit from Qiagen according to the manufacturer´s instructions. Viral DNA presence was analyzed by qPCR (Applied Biosystem 7500) with Takyon^TM^ Low Rox Probe MasterMix dTTP Blue (Eurogentec) using primers detecting gD HSV-1 sequence [45] (Forward: 5’-CGGCCGTGTGACACTATCG-3’, Reverse:5’-CTCGTAAAATGGCCCCTCC-3’, Probe: FAM-CCATACCGACCACACCGACGAACC-TAMRA) and primer pair specific for GAPDH (Forward: 5’-CGCTCTCTGCTCCTCCTGTT-3’, Reverse:5’-CCATGGTGTCTGAGCGATGT-3’, Probe: 5’-CAAGCTTCCCGTTCTCAGCC-3’). qPCR was performed using as template the supernatant media collected from infected plates and boiled for 5 min at 95°C. Serial dilutions of known-titre <u>HSV1</u> solution were used as a standard curve to extrapolate the genomes/ml in the samples. Dilutions series of known standards were analysed in each plate. Standard curve was created comparing the known copies/reaction value with the mean of the Crossing threshold value (Ct value). Trendline was extrapolated by software, used to interpolate the genomes/ml of the analysed sample and data were compared with the control. When DNA extracted from pellet was analysed, both sets of primers where used in order to assess the cellularity of the sample. Values were calculated with ΔΔCT analysis and data are expressed in relative genome copy compare to the control.

## Results

### Isolation of cellular proteins involved in primary envelopment

During nuclear egress virus particles first acquire a primary envelope from the INM and then this fuses with the ONM to release the particles into the cytoplasm. Little thought has gone into what happens to the membrane and membrane proteins involved in this process after this point, but both proteins in the primary envelope and those involved in fusion with the ONM should be released into the ONM. Due to the continuity of the ONM with the ER such proteins are as likely to diffuse throughout the ER as to re-translocate back to the INM (Fig. 1A). An ER-enriched membrane fraction can be obtained by preparation of microsomes, where cellular membranes are separated by floating through sucrose after removal of intact nuclei and mitochondria [46, 47]. Therefore to determine potential host proteins involved in herpesvirus nuclear egress, we performed a proteomic analysis on isolated ER-enriched microsomal membrane (MM) fractions from HSV-1 and mock infected cells, searching for NE proteins with increased abundance in the ER during viral infection.

**Figure 1.**
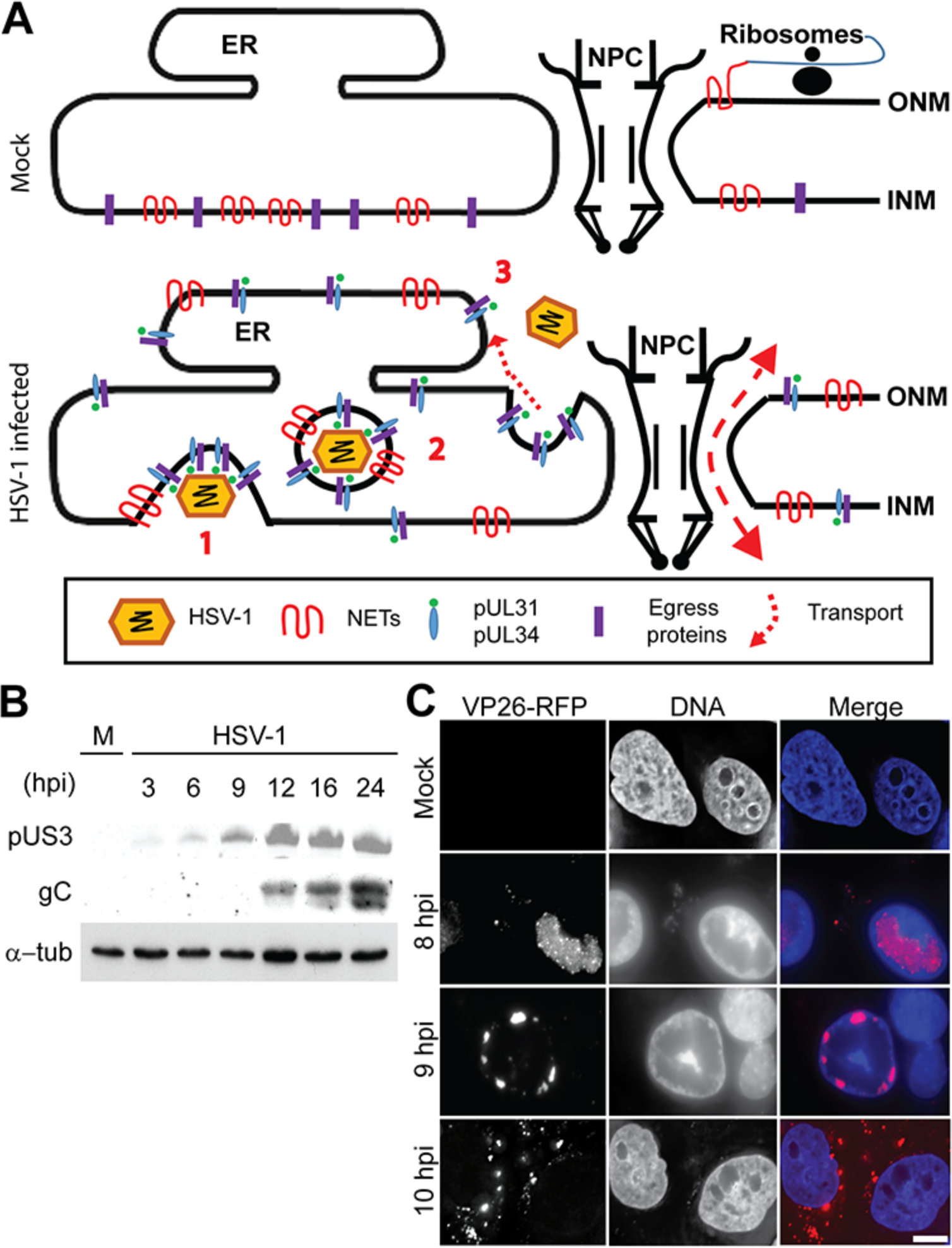
Optimization of conditions for isolation of HSV-1 infected MMs. (**A**) Schematic representation of the NE in mock-infected (top diagram) and HSV-1 infected (bottom diagram) cells. NE transmembrane proteins (NETs) and potentially as yet unidentified egress proteins are only transiently in the ER and ONM after synthesis (top diagram). They diffuse laterally through the peripheral channels of the NPC to reach the INM. In HSV-1 infected cells, nucleocapsids bud at the INM (1) into the lumen followed by scission of the INM-derived vesicle to form primary virions (2). During de-envelopment, the INM-derived primary envelope fuses with the ONM, releasing proteins involved in primary envelopment (3). Due to continuity of the ONM with the ER these proteins can diffuse into the ER or re-translocate back to the INM. (**B**) An expression timecourse of US3 and gC viral proteins was analyzed by Western blot to determine the optimal time to isolate HSV-1 infected MMs before significant secondary envelopment has occurred. Cell lysates from Hela cells infected at MOI 10 were prepared at indicated times post-infection. α-tubulin was used as a loading control. (**C**) Fluorescence microscopy images of an HSV-1 strain with VP26 capsid protein fused with RFP in Hela cells after 8, 9 and 10 hpi. At 8 hpi most of the VP26 signal was located inside the nucleus of infected cells. At 9 hpi cells started to show perinuclear localization of VP26. At 10 hpi some viral particles were detected in the cytoplasm of infected cells. Scale bar, 10 μm.

As proteins involved in secondary envelopment might also accumulate in the MM fraction, it was necessary to first determine the optimal time during infection where nuclear egress has started before secondary envelopment. To do so two viral proteins, pUS3 and glycoprotein C (gC), were tracked for expression during the course of infection in the HeLa cells used. pUS3 is a serine/threonine viral kinase thought to support fusion of primary enveloped particles with the ONM because its deletion results in accumulation of particles in the NE and ER lumen [15, 16]. It is expressed before primary envelopment/de-envelopment, peaking around the time that secondary enveloped particles are first detected. In contrast, gC is a component of the final envelope that is acquired in the Golgi when viral particles have accumulated in the cytoplasm after their release from the nucleus. In the HeLa cells pUS3 was detectable at 3 h post-infection (hpi), but did not begin to be highly expressed until 9 hpi and reached maximal levels by 12 hpi (Fig. 1B). At this time, gC was only becoming detectable and increased in levels over time to peak at 24 hpi. Based on this result, it was determined that to minimize capturing changes in the ER and Golgi associated with secondary envelopment in the MM fractions the cells should be isolated before 12 hpi.

A second analysis determined the timing when many viral particles could be detected in the nucleus, but very few had entered the cytoplasm. An HSV-1 strain with capsid protein VP26 tagged with RFP was followed in a time course by fluorescence microscopy to reveal that at 8 hpi punctate spots indicative of viral particles are inside the nuclei in most of infected cells while at 10 hpi some viral particles are already visible in the cytoplasm (Fig. 1C). Therefore, to limit changes associated with secondary envelopment the isolation of ER-enriched MMs was performed between 8 and 9 hpi.

HeLa cells were infected at a multiplicity of infection (MOI) of 10 in order to ensure a rapid and even infection throughout the cell population. At 8 hpi 30 L of cells were collected from roller bottles by trypsinization and collected by centrifugation. MMs and NEs were prepared according to published protocols [31, 32] (Fig. 2A). Briefly, cells were hypotonically lysed with Dounce homogenization (Fig. 2B) and intact nuclei were removed by centrifugation. Nuclei were further processed to prepare NEs by further douncing with a tight-fitting homogenizer, pelleting the intact nuclei through sucrose to remove other membranes and contaminants, then digesting and washing away chromatin. To prepare MMs, the post-nuclear supernatant was first depleted of mitochondria by centrifugation and then floated through sucrose layers where the MMs accumulate just above the 1.9 M sucrose layer. As expected, the NE and MMs fractions were clearly distinct in composition (Fig. 2C) and Western blotting for NE and ER markers revealed the expected distributions for the fractions (Fig. 2D). NE markers Lamin A/C and LAP2β were only found in the NE fraction while the ER marker calnexin was predominantly in the MM fraction with a small amount in the NE fraction, as expected since the ONM is not only continuous with the ER but also studded with ribosomes and many other ER proteins. Importantly, both NE and MM fractions were free of contamination with the Golgi membranes that function in secondary envelopment as the Golgi marker GM-130 was absent from the fractions (Fig. 2E).

**Figure 2.**
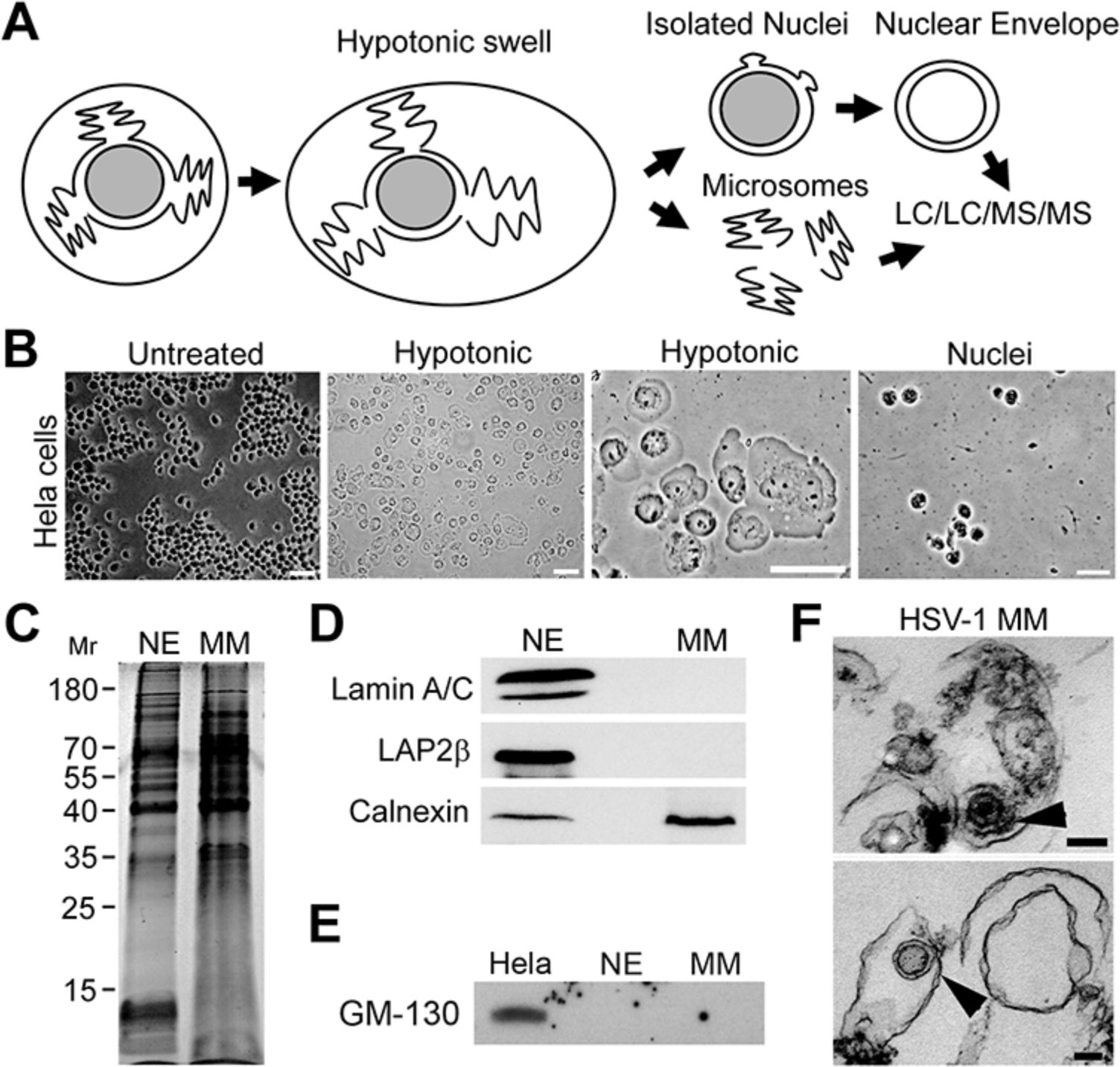
Isolation of MMs from HSV-1 infected cells to identify host proteins involved in HSV-1 primary envelopment. (**A**) Method schematic. Cells were lysed after hypotonic swelling and crude nuclei separated from crude cellular membranes by pelleting. Nuclei were subsequently cleaned of contaminating membranes by pelleting through sucrose gradients. NEs were prepared by subsequently digesting and removing DNA and RNA. Separately, MMs were prepared from the crude cellular membrane fraction by floating (after pelleting mitochondria) through sucrose gradients and taking the microsome-enriched fraction. Cellular fractions from mock-infected and HSV-1 infected cells were analyzed by LC/LC/MS/MS to identify host proteins diffusing from the INM into the ONM and ER during primary envelopment.(**B**) Cell fractions. Mock-infected and HSV-1 infected Hela cells were swollen hypotonically (middle panels) and dounce homogenized (right panel) to release nuclei. Phase-contrast light microscope images are shown. Scale bars, 10 μm. (**C**) Coomassie-stained gel of NE and MMs fractions showed a clear difference in protein composition. (**D**) Western blot of fractions from (C) stained with ER and NE markers to determine fraction purity. The ER marker calnexin was present in both NE and MM fractions as expected because the ONM is continuous with the ER and many proteins are shared. In contrast, the NE markers lamin A/C and Lap2β were absent from MMs. Similar amounts of total protein were loaded as in (C). (**E**) To test for possible Golgi contamination in the preps, the NE and MM fractions were blotted for Golgi marker GM-130. Equal protein loading from HeLa cells confirmed that the antibody was working. (**F**) Ultrastructure of isolated MMs from HSV-1 infected Hela cells. Electron micrographs showed the characteristic single-membrane structure of the MMs. Arrows point to electron dense symmetrical structures of around 100 nm diameter that probably represent primary viral particles with the bottom one likely lacking a packaged genome. Scale bars, 100 nm.

Electron microscopy of the isolated HSV-1 MMs revealed the expected appearance of membrane vesicles and, in addition, electron dense symmetrical structures of ~100 nm diameter with the appearance of virus particles were sometimes observed within the vesicles (Fig. 2F). These might have been produced during douncing due to the weakened NE. These particles were not immunogold labeled to confirm their identity, but may be primary enveloped particles. This suggests that this approach, in conjunction with immunogold labeling, might be modified in the future to obtain a pure population of primary enveloped particles.

### Vesicle fusion proteins identified in HSV-1 infected MMs

MMs from HSV-1 infected cells and NEs and MMs from mock-infected cells were analyzed by mass spectrometry using the Multi-dimensional Protein Identification Technology (MudPIT) LC/LC/MS/MS approach [34, 35]. As expected from the electron microscopy structures with the appearance of viral particles, many HSV-1 proteins were identified specifically in the infected MMs. These are most likely primary enveloped virions as all but pUL17 were detected out of 8 viral proteins previously found in an attempt to isolate and identify the composition of primary enveloped virions [48] (Table 1). This list included several known tegument proteins, the pUL34 protein already known to be important for nuclear egress and glycoprotein D, indicating that some proteins in the final envelope also participate in primary envelopment.

**Table 1.**
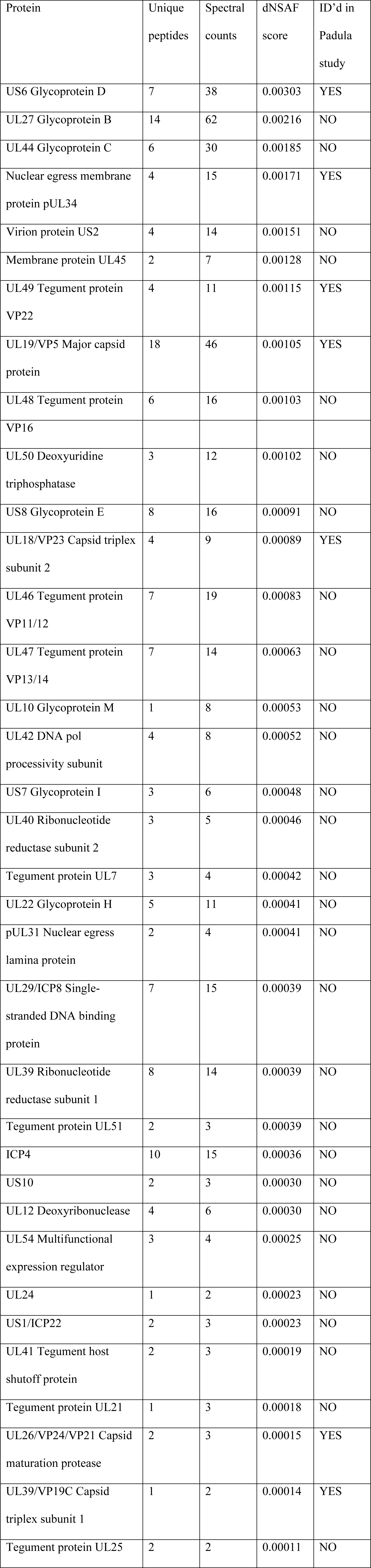
Abundance of viral proteins identified in HSV-1 infected MMs.

| Protein | Unique peptides | Spectral counts | dNSAF score | ID'd in Padula study |
| --- | --- | --- | --- | --- |
| US6 Glycoprotein D | 7 | 38 | 0.00303 | YES |
| UL27 Glycoprotein B | 14 | 62 | 0.00216 | NO |
| UL44 Glycoprotein C | 6 | 30 | 0.00185 | NO |
| Nuclear egress membrane protein pUL34 | 4 | 15 | 0.00171 | YES |
| Virion protein US2 | 4 | 14 | 0.00151 | NO |
| Membrane protein UL45 | 2 | 7 | 0.00128 | NO |
| UL49 Tegument protein VP22 | 4 | 11 | 0.00115 | YES |
| UL19/VP5 Major capsid protein | 18 | 46 | 0.00105 | YES |
| UL48 Tegument protein | 6 | 16 | 0.00103 | NO |
| VP16 |  |  |  |  |
| UL50 Deoxyuridine triphosphatase | 3 | 12 | 0.00102 | NO |
| US8 Glycoprotein E | 8 | 16 | 0.00091 | NO |
| UL18/VP23 Capsid triplex subunit 2 | 4 | 9 | 0.00089 | YES |
| UL46 Tegument protein VP11/12 | 7 | 19 | 0.00083 | NO |
| UL47 Tegument protein VP13/14 | 7 | 14 | 0.00063 | NO |
| UL10 Glycoprotein M | 1 | 8 | 0.00053 | NO |
| UL42 DNA pol processivity subunit | 4 | 8 | 0.00052 | NO |
| US7 Glycoprotein I | 3 | 6 | 0.00048 | NO |
| UL40 Ribonucleotide reductase subunit 2 | 3 | 5 | 0.00046 | NO |
| Tegument protein UL7 | 3 | 4 | 0.00042 | NO |
| UL22 Glycoprotein H | 5 | 11 | 0.00041 | NO |
| pUL31 Nuclear egress lamina protein | 2 | 4 | 0.00041 | NO |
| UL29/ICP8 Single-stranded DNA binding protein | 7 | 15 | 0.00039 | NO |
| UL39 Ribonucleotide reductase subunit 1 | 8 | 14 | 0.00039 | NO |
| Tegument protein UL51 | 2 | 3 | 0.00039 | NO |
| ICP4 | 10 | 15 | 0.00036 | NO |
| US10 | 2 | 3 | 0.00030 | NO |
| UL12 Deoxyribonuclease | 4 | 6 | 0.00030 | NO |
| UL54 Multifunctional<br>expression regulator | 3 | 4 | 0.00025 | NO |
| UL24 | 1 | 2 | 0.00023 | NO |
| US1/ICP22 | 2 | 3 | 0.00023 | NO |
| UL41 Tegument host<br>shutoff protein | 2 | 3 | 0.00019 | NO |
| Tegument protein UL21 | 1 | 3 | 0.00018 | NO |
| UL26/VP24/VP21 Capsid<br>maturation protease | 2 | 3 | 0.00015 | YES |
| UL39/VP19C Capsid<br>triplex subunit 1 | 1 | 2 | 0.00014 | YES |
| Tegument protein UL25 | 2 | 2 | 0.00011 | NO |

Interestingly, this earlier study failed to identify pUL31, which is known to also be important for primary envelopment and forms a complex with pUL34. In our dataset we found both pUL31 and pUL34 (Table 1). Additionally, we found several other tegument proteins and glycoproteins; however, the spectral abundance of the glycoproteins broke down into two groups. The first group including gD, gB and gC were all based on abundance estimated from spectral counts present at roughly 6x greater abundance than the others detected, gE, gM, gI and gH (Table 1). Association of gD, gB and gC with HSV-1 primary virions have been suggested in different studies [49, 50] and gB has been implicated in fusion with the ONM during egress [17, 18]. This might suggest that the more abundant set participate in primary envelopes and nuclear egress while the less abundant glycoproteins are simply beginning to be expressed and in the process of moving through the ER to the Golgi to support subsequent secondary envelopment. Nonetheless, though the trend may be real, it is likely an oversimplification as gH was also implicated in ONM fusion [17, 18] despite that our data place it in the less abundant set. We postulate that the additional tegument proteins are likely to occur in primary enveloped particles because, not being transmembrane, they would likely have been lost during the MM purification unless they are part of primary enveloped virions captured in the MMs (Fig. 2F).

We postulated that host cell proteins from the INM either captured during the process of primary envelopment/ de-envelopment or actively participating in this process would be released into the ONM and diffuse into the ER (Fig. 1A-bottom). Therefore, one would expect the proteins of greatest interest to occur in the NE fraction and to be increased in the HSV-1 infected MM fraction over the mock-infected MM fraction. Thus, we extracted all proteins in the HSV-1 infected MMs that were at least 30% increased over the mock-infected MMs based on their dNSAF values, a measure of the percentage of the total mass of the mass spectrometry run accounted for by a particular protein based on spectral abundance and its molecular weight (see Materials and Methods), and occurred also in the NE fraction. These proteins were analyzed for specific gene ontology (GO)-terms and plotted as a percentage of the dNSAF obtained for each protein summed for a particular GO term. This was then compared to the distribution of specific GO-terms for the whole genome (Fig. 3A). Several GO terms were strongly enriched in the select HSV-1 infected MMs including protein phosphorylation, membrane organization and nucleocytoplasmic transport. However, the most striking both for its >3-fold increase and for its potential relevance to herpesvirus nuclear egress was the vesicle fusion protein category.

**Figure 3.**
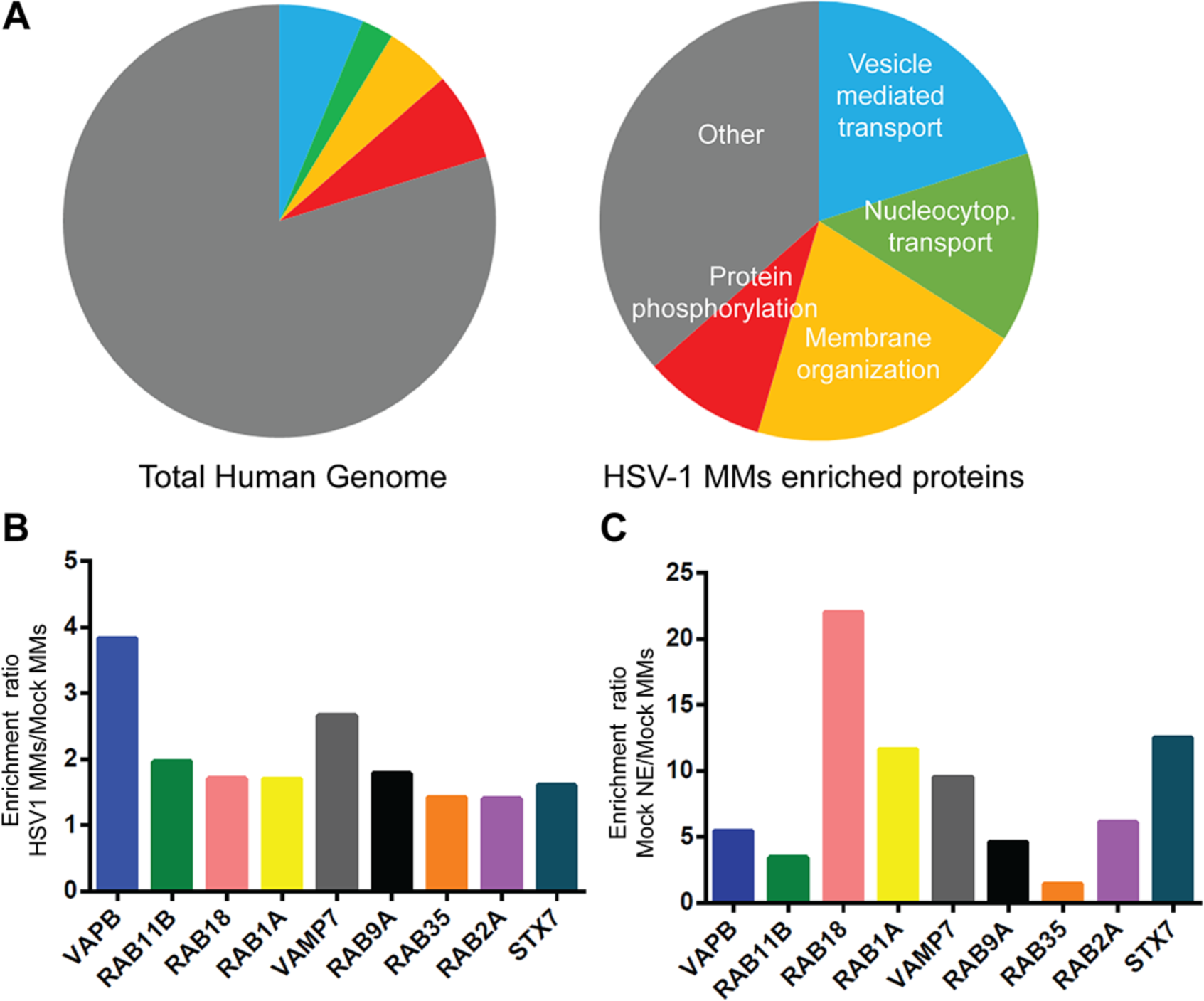
Enrichment of vesicle fusion proteins in HSV-1 infected MMs identified by mass spectrometry. (**A**) Gene ontology (GO) classification for enriched proteins from HSV-1 infected MMs compared with their representation in the total human genome. While the Vesicle mediated transport GO-category reflects only 6% of genes encoded in the human genome, it represents 20% of the enriched proteins detected in HSV-1 MMs. Other interesting categories were proteins involved in nucleocytoplasmic transport (14% in HSV-1 MMs versus the 2.4 % in total human genome) and membrane organization proteins (20.5% in HSV-1 MMs versus 4.9% in total human genome). (**B and C**) Protein abundance estimates based on spectral counts and protein molecular weight as a percentage of the total mass of spectra identified per mass spectrometry run (dNSAF, see Methods section) were used to calculate the relative enrichment between the three mass spectrometry datasets. The ratio of dNSAF values for different vesicle fusion proteins between HSV-1 infected MMs and mock-infected MMs (**B**) and between NEs and mock-infected MMs (**C**) are shown. Only those vesicle fusion proteins that had a HSV-1 infected MMs:mock-infected MMs ratio of >1.3 are presented.

The stringency of the selection criteria was then increased for this set of vesicle fusion proteins, only considering those that had at least 5 spectra in the NE fraction to suggest that they are reasonably abundant and that also were at least 1.3-fold more abundant based on spectral counts in the HSV-1 infected MMs over the mock-infected MMs. The relative enrichment in each of the fractions for each of these select candidates was plotted (Fig. 3B,C) and their peptide and spectra data are given in Table 2. Three proteins were chosen for detailed analysis: VAPB because it was the most enriched in the HSV-1 infected MMs over the mock-infected MMs, Rab18 because it was the most enriched in NEs over mock-infected MMs, and Rab11b because it was roughly in the middle.

**Table 2.**
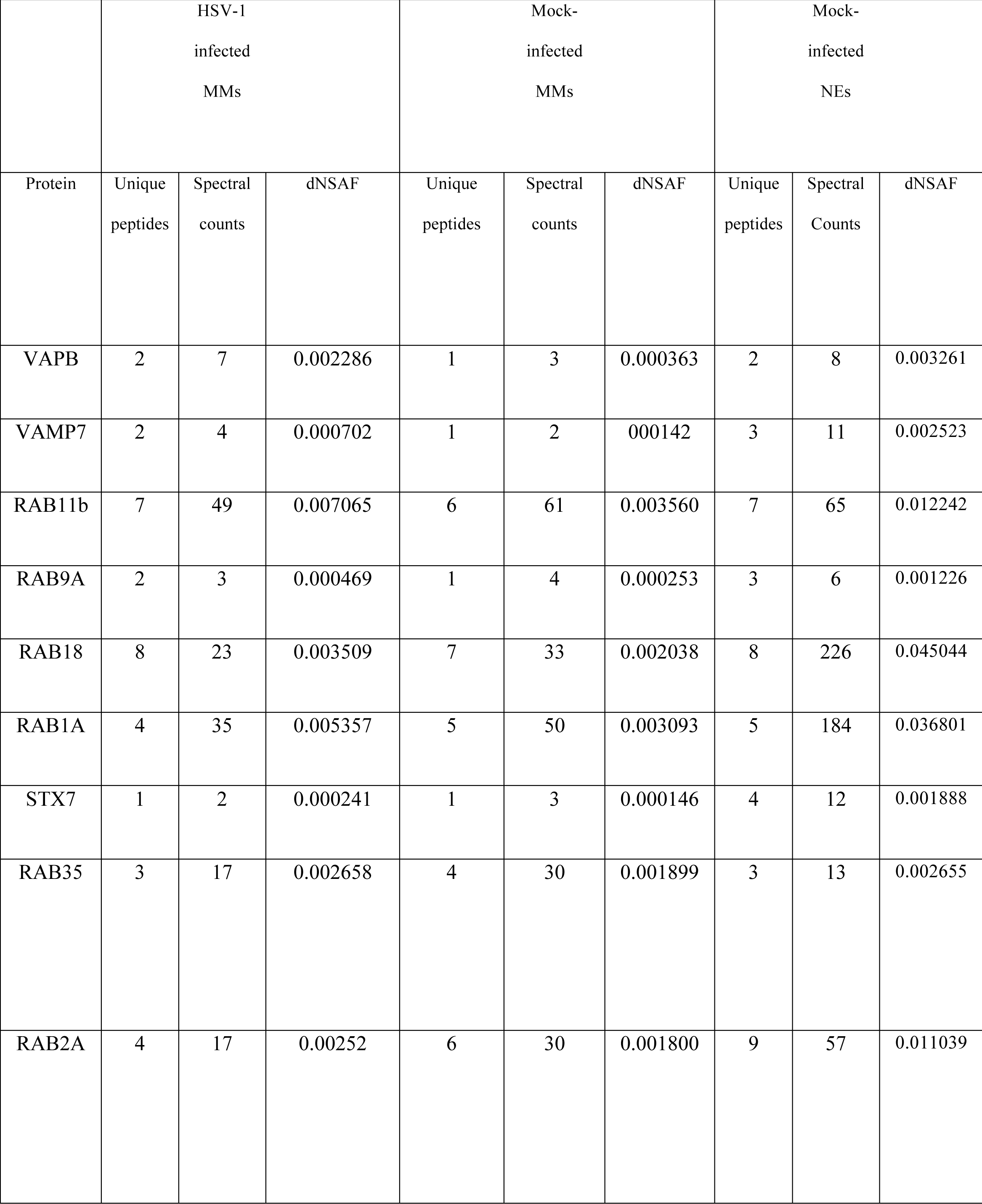
Abundance of vesicle fusion proteins enriched in HSV-1 infected MMs.

|  | HSV-1<br>infected<br>MMs |  |  | Mock-infected<br>MMs |  |  | Mock-infected<br>NEs |  |  |
| --- | --- | --- | --- | --- | --- | --- | --- | --- | --- |
| Protein | Unique<br>peptides | Spectral<br>counts | dNSAF | Unique<br>peptides | Spectral<br>counts | dNSAF | Unique<br>peptides | Spectral<br>Counts | dNSAF |
| VAPB | 2 | 7 | 0.002286 | 1 | 3 | 0.000363 | 2 | 8 | 0.003261 |
| VAMP7 | 2 | 4 | 0.000702 | 1 | 2 | 0.000142 | 3 | 11 | 0.002523 |
| RAB11b | 7 | 49 | 0.007065 | 6 | 61 | 0.003560 | 7 | 65 | 0.012242 |
| RAB9A | 2 | 3 | 0.000469 | 1 | 4 | 0.000253 | 3 | 6 | 0.001226 |
| RAB18 | 8 | 23 | 0.003509 | 7 | 33 | 0.002038 | 8 | 226 | 0.045044 |
| RAB1A | 4 | 35 | 0.005357 | 5 | 50 | 0.003093 | 5 | 184 | 0.036801 |
| STX7 | 1 | 2 | 0.000241 | 1 | 3 | 0.000146 | 4 | 12 | 0.001888 |
| RAB35 | 3 | 17 | 0.002658 | 4 | 30 | 0.001899 | 3 | 13 | 0.002655 |
| RAB2A | 4 | 17 | 0.00252 | 6 | 30 | 0.001800 | 9 | 57 | 0.011039 |

### Knockdown of vesicle fusion proteins results in nuclear accumulation of virus particles and significant reduction of HSV-1 viral titers

The proteins redistributed according to the mass spectrometry data could directly benefit the virus or reflect a general perturbation of vesicle trafficking during infection. To determine if they benefit the virus, siRNA was used to knock down VAPB, Rab18 and Rab11b and the effect on viral titers tested. Rab24 and Rab1a were also knocked down as controls. Rab24 was used as a negative control as it did not increase in the HSV-1 infected MMs compared to mock-infected MMs. Rab1a was used as a positive control because this protein required for ER-to Golgi complex transport [51] has been shown to be involved in HSV-1 mature particle assembly (secondary envelopment) and its knockdown reduces viral growth by 60% [52].

HeLa cells were transfected with siRNA oligos and after 48 h the cells were lysed and analyzed by Western blot (Fig. 4A). VAPB was knocked down to nearly undetectable levels while the Rab11b and Rab18 level was knocked down by at least 85%. The controls Rab1a and Rab24 were knocked down similarly to roughly 80%.

**Figure 4.**
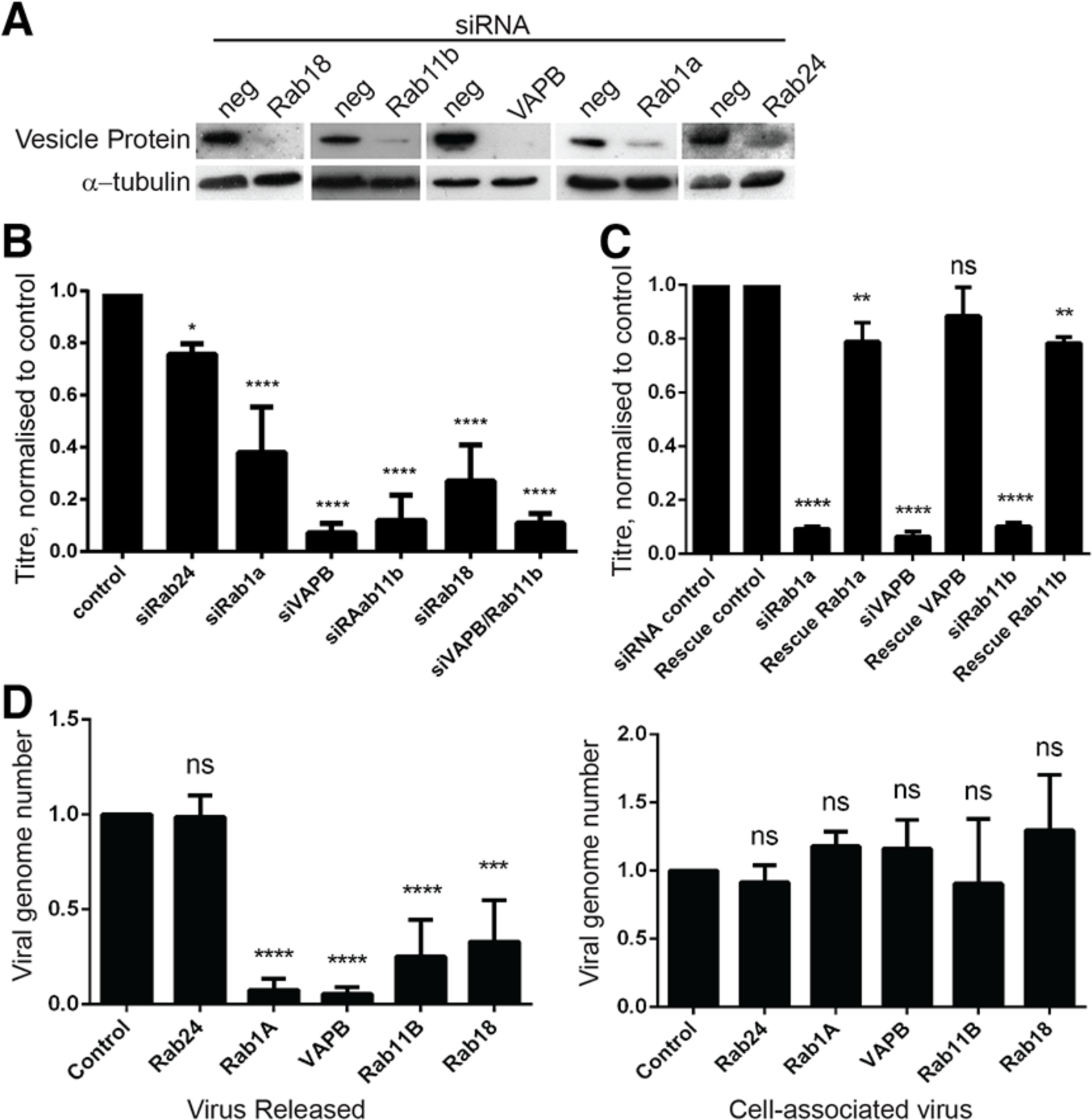
VAPB, Rab11b and Rab18 knockout inhibits HSV-1 infection. (**A**) HeLa cells were transfected with siRNA oligos for the indicated proteins and after 48 h the cells were lysed and analyzed by Western blot with the indicated antibodies to check protein knockdown. (**B**) Hela cells were transfected with the control, Rab24, Rab1a, VAPB, Rab11b, Rab18, or a combined VAPB and Rab11b siRNA. After 48 h, cells were infected with HSV-1 (MOI, 10) and supernatant virions were collected at 16 hpi. Titers were established on U2OS cells and bars represent the average of three independent experiments, normalized to the control siRNA. Error bars are to the standard deviation and the p-values for comparing each condition with the control are shown (****p ≤ 0.0001; * p≤0.005). (**C**) Rescue experiments with cells expressing wild-type protein resistant to the siRNAs were performed. In all cases the cells carrying both the rescue plasmid and the siRNAs recovered to roughly 80-90% of the control virus titers. (**D**) HSV-1 genome copy numbers were determined from released viral particles and from cell pellets by qPCR. HSV-1 genome copy number from viral particles released from infected cells is reduced by 87% in siRab11b, 80% in siRab18 HSV-1 and 90% in siVAPB. However, HSV-1 genome copy number is unaffected in cell-associated virus compared with control. The p-values for comparing each condition with the control indicate a statistical significance in released virus while there is no significant difference in cell-associated virus (ns). All statistical tests were performed by one-way ANOVA and multiple comparisons were done by Dunnett´s test.

Another set of siRNA transfections was undertaken and at 48 h infected cells were subsequently infected with HSV-1 at a MOI of 10. At 16 hpi supernatant from these cells was collected and tested by virus titration assay. As expected by previous studies, very little effect of Rab24 knockdown could be observed [53], while Rab1a knock down reduced viral titers by 62% (Fig. 4B). Compellingly, all three vesicle-fusion proteins identified by our proteomic approach exhibited a stronger reduction in virus titers than the Rab1a knockdown. As an average of three separate experiments, Rab18 knockdown yielded ~70% reduction in virus titers (Fig. 4B). Rab11b knockdown had an even stronger effect with ~85% reduction in virus titers. Finally, VAPB knockdown yielded over 90% reduction in titer. These were not off-target effects of the siRNAs as rescue experiments yielded full recovery of virus titers (Fig. 4C).

If the vesicle fusion proteins function in different complementing pathways, then combinatorial knockdown might be expected to yield a stronger effect. However, when VAPB and RAB11b were depleted together, the resultant titer was no greater than for individual knockdowns (Fig. 4B, rightmost bar). To determine if indirect effects of vesicle trafficking on viral replication might underlie the reduction in virus titers, the effect of VAPB, Rab11b and Rab18 on viral genomes produced was also tested. Supernatants and cell pellets were collected and analyzed by qPCR. As expected from the virus titer experiments, the knockdowns resulted in a very significant reduction in virion (as measured by viral DNA copy number) release from the cell with the control being unaffected, similar to what was found in the titration data (Fig. 4D, left graph). Although it would have been ideal to isolate cytoplasmic vs nuclear preparations, they could not be prepared due to nuclear weakness and breakage in the infected cells. However, qPCR for viral genomes in the infected whole cell pellets revealed that virus production was not significantly altered between the control and VAPB, Rab11b or Rab18 knockdowns (Fig. 4D, right graph). In fact, viral genome numbers in the siRNA-treated infected cells were all slightly higher than the control, presumably because of their accumulation when the egress pathways are disrupted. Together the reduction in excreted HSV-1 genomes despite their effective production in the nucleus is consistent with a role of these vesicle fusion proteins in steps for egress as opposed to replication.

These results indicate that VAPB, Rab18 and Rab11b all play important roles in viral maturation, though they do not distinguish whether the loss in titers reflects a function in nuclear egress or secondary envelopment. To distinguish these possibilities, we tested for virus accumulation in the nucleus as this would only be observed if nuclear egress is affected. Electron microscopy of control cells infected with virus yielded the expected distribution with some assembling nuclear particles and many virus particles accumulating in the cytoplasm at 16 hpi (Fig. 5, upper left panel). The same was observed for the Rab24 knockdown infected cells (Fig. 5, upper right panel). However, the knockdowns for Rab11b, Rab18 and VAPB all revealed accumulation of a much larger number of virus particles in the nucleus with very little detectable in the cytoplasm at 16 hpi (Fig. 5, other image panels). For VAPB in particular some images revealed enveloped particles in lumenal extensions of the NE (Fig. 5B). Counting of nuclear and cytoplasmic particles in several images (Fig. 5C) was consistent with the visually observed tendency to accumulate particles in the nucleus; however, as electron microscopy only captures those particles in a particular section of the nucleus we sought better quantification using fluorescence in situ hybridization (FISH).

**Figure 5.**
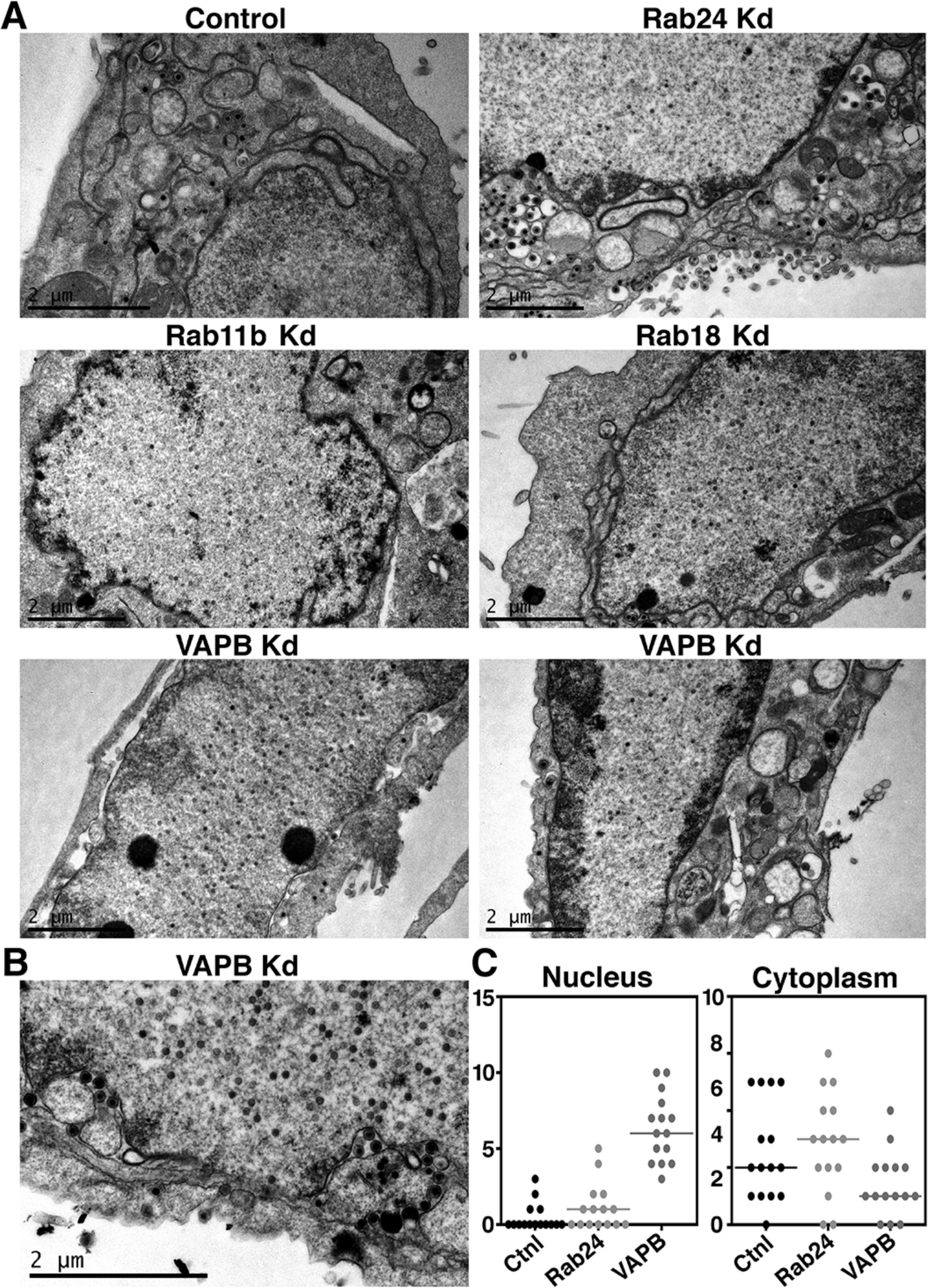
Electron microscopy reveals accumulation of virus particles in the nucleus with vesicle protein knockdowns. In control and Rab24 knockdown cells some non-enveloped virus particles could be observed in the nucleoplasm, but many both enveloped and non-enveloped particles could also be observed in the cytoplasm as well as released mature particles just outside the cell. In contrast many more nucleoplasmic non-enveloped particles were observed in the Rab11b, Rab18 and VAPB knockdowns. Very few enveloped particles were observed for these three knockdowns in either the cytoplasm or nucleus, but some seemingly enveloped particles could be seen for Rab11b and VAPB knockdowns in association with the nuclear envelope and non-enveloped particles for the Rab18 knockdown. Bottom right panel. Quantification of particles for the control and Rab24 knockdowns revealed relatively few particles in the nucleus compared to the cytoplasm whereas for the VAPB knockdown there were mostly particles in the nucleus and few in the cytoplasm. Further quantification was not engaged because this was done more accurately by fluorescence in situ hybridization (see Figure 6).

Infected cells with control and vesicle fusion protein knockdowns were hybridized with a biotin-labeled probe for the gene encoding the HSV-1 ICP27 protein and subsequently visualized by incubation with streptavidin conjugated fluorophore. The cells were pre-treated with RNase so that ICP27 transcripts would not be recognized, but only viral genomes. Co-staining with DAPI identified the nuclear boundaries and imaging revealed virus accumulating in the cytoplasm in the non-target siRNA control cells, the Rab24 knockdown that had not changed according to the mass spectrometry results, and the Rab1A knockdown that is known to affect secondary envelopment but not nuclear egress (Fig. 6A). In contrast, the FISH signal for viral genomes was visually restricted to the nucleus in the VAPB, Rab11b and Rab18 knockdowns (Fig. 6A). The intensity of total FISH signal in each cell and also that just in the nucleus using the DAPI staining as a mask was determined. The total signal divided by the nuclear signal was then plotted so that the amount of signal outside the nucleus is reflected in values above 1 (Fig. 6B). All three controls exhibited a clear increase in cytoplasmic virus while VAPB, Rab11b and Rab18 knockdowns all remained around 1. Therefore, these vesicle fusion proteins clearly act at the level of nuclear egress.

**Figure 6.**
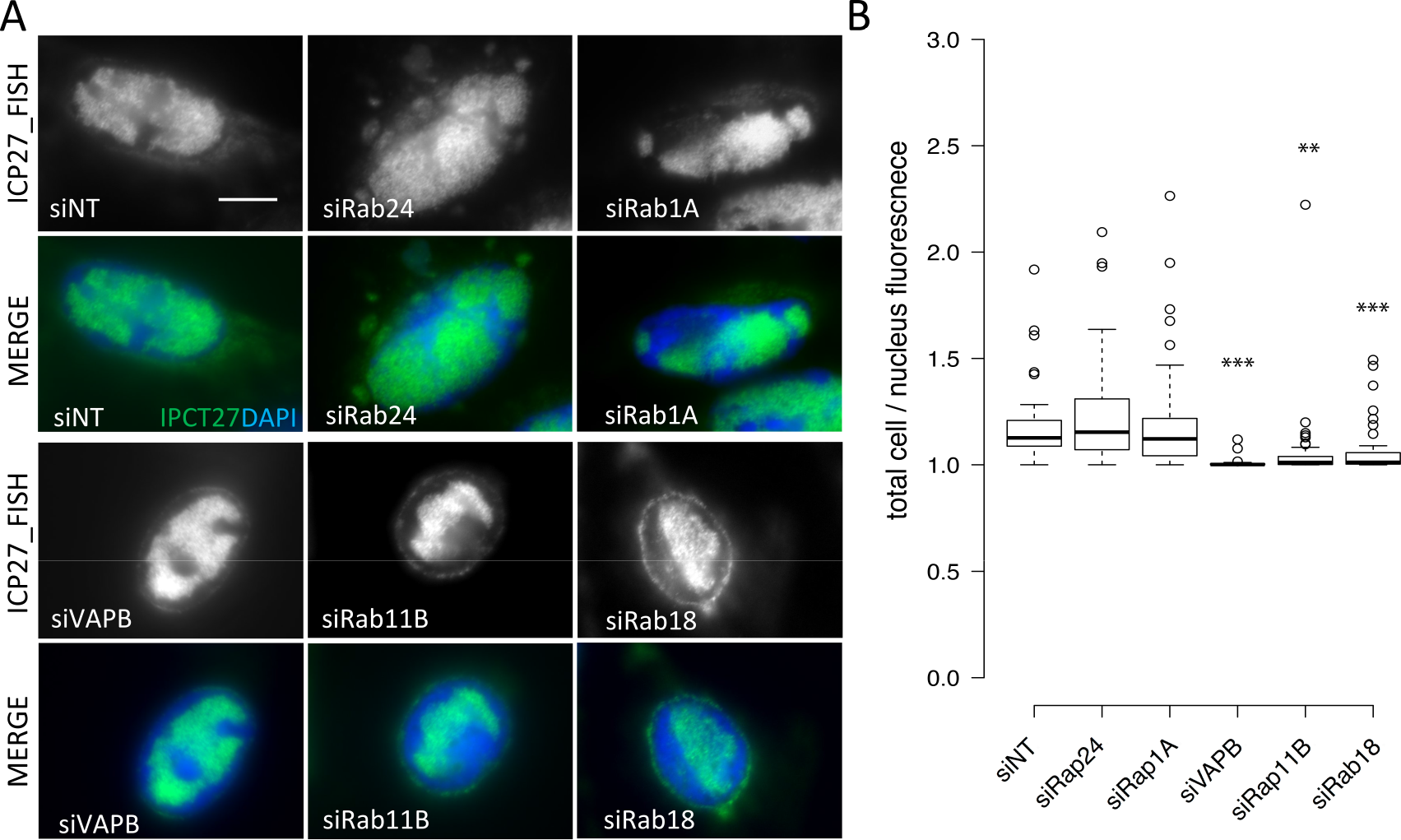
Fluorescence in situ hybridization (FISH) to quantify nuclear and cytoplasmic virus particles in vesicle fusion protein knockdown cells. (**A**) The virus gene ICP27 was used as a probe and labeled with biotin. Cells were knocked down for vesicle fusion proteins as in Figures 4 and 5, infected with HSV-1 and at 16 hpi fixed and processed for FISH. The hybridized virus gene was visualized with streptavidin conjugated to Alexa488 dye and imaged by immunofluorescence microscopy. Cells were co-stained with DAPI to identify the nucleus. Scale bar, 10 μm. (**B**) Using the DAPI nuclear staining to generate a mask of the nuclear area the nuclear pools of hybridized virus ICP27 DNA were quantified. The total hybridized ICP27 DNA in the same cell was also quantified and plotted divided by the nuclear signal so that values above 1 reflect the cytoplasmic pool of viral genomes. A clear increase in cytoplasmic viral genomes can be seen for the non-target siRNA control, the Rab24 and the Rab1A knockdowns while no notable increase in cytoplasmic viral genomes was observed for VAPB, Rab11b and Rab18 knockdowns. Statistical measurements were performed using a 2-tailed ANOVA analysis: ***p<0.001.

### VAPB is recruited to the NE during HSV-1 infection

We next sought to explore the intracellular distribution of these proteins during HSV-1 infection. As rabbit antibodies often react with Fc receptors expressed by HSV-1, we were only able to test the distribution of VAPB for which we were able to obtain a good mouse monoclonal antibody. HeLa cells were infected and fixed and stained at different stages of infection (8 and 16 hpi), followed by indirect immunofluorescence analysis to test for confirmation of the VAPB NE localization indicated by the proteomics data (Fig. 7A). In order to select for analysis only those cells infected by HSV-1, the infection was performed using an HSV-1 strain in which ICP27 was tagged with GFP. VAPB was visibly observed to accumulate at the NE over time in the infected cells compared to the mock-infected cells with the strength of the signal dissipating through the ER in keeping with the proteomics results. This re-localization of VAPB was observed in three independent experiments and when NE intensities were quantified and compared with ER intensities, it clearly demonstrated a statistically significant increase in detection of these fusion proteins at the NE prior to the initiation of nuclear egress at 8 hpi. To quantify the relative signal for VAPB in the NE and ER, multiple lines were drawn through the middle of the nucleus of at least 40 cells and the signal intensity at the edge of the nucleus based on DAPI staining for the DNA was measured and also the signal in the ER along the same line but 2 μm away from the nucleus was also measured. The ratio of NE to ER signal intensity was plotted (Fig. 7A lower graph panel), revealing a strongly statistically significant near doubling of the NE:ER signal ratio between the mock-infected cells and the 16 hpi timepoint with the 8 hpi timepoint in between.

**Figure 7.**
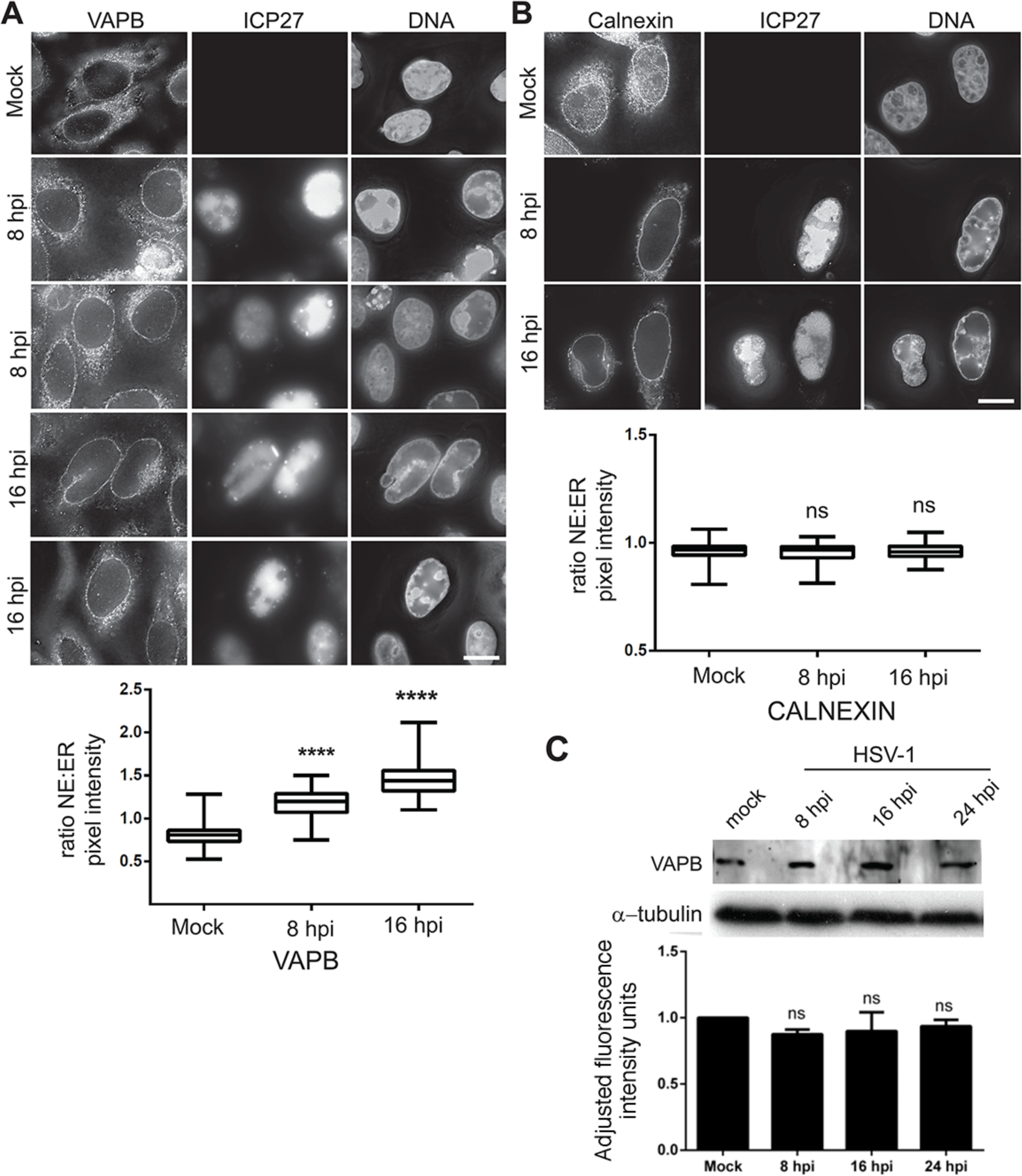
VAPB is recruited to the NE during HSV-1 infection. (**A**) Hela cells were either mock infected or infected with MOI 10 HSV-1 vBSGFP27 (encoding WT ICP27 fused to GFP). At 8 and 16 hpi cells were fixed and processed for immunofluorescence microscopy with VAPB mouse monoclonal antibodies and DNA stained with DAPI. ICP27-GFP is shown to identify infected cells. Scale bar, 10 μm. The VAPB signal appeared visually to increase at the NE during infection. To quantify this putative NE redistribution, the relative pixel intensities in the ER and NE were quantified. For each cell five lines were drawn through the middle of the nucleus and pixel intensity was measured at a point in the nuclear rim (based on DAPI staining) and at a point 2 μm distant into the ER and the NE/ER ratio was calculated. Boxplots from 30 cells are shown in the graph below the images with the median (central line) and the error bars (grey) marked. Each sample was compared with its control (mock cells) by one-way ANOVA analysis followed by Holm-Sidak´s multiple-comparison test. Significant p-values (****p ≤ 0.0001) illustrate the general trend of these vesicle fusion proteins to accumulate at the NE upon infection. (**B**) As a control the same analysis was performed staining for the ER and ONM marker Calnexin. In this case no significant change in the relative distribution was observed. (C) Analysis of total protein levels of VAPB in HSV-1 infected cells during infection. Lysates were prepared from mock-infected HeLa cells or HeLa cells infected with WT HSV-1 with MOI 10 after 8, 16 and 24 hpi in three separate experiments. Each lysate was analyzed by Western blots with the antibodies shown and using fluorophore-conjugated secondary antibodies for quantification by Li-COR. No obvious differences were observed in VAPB protein levels at any of the time points.

In contrast, staining for calnexin yielded no notable visible increase at the NE in infected cells and quantification confirmed that there was no increase in the NE:ER ratio (Fig. 7B). The VAPB increase at the NE during HSV-1 infection is not due to changes in the protein levels because total cell lysates analyzed by Western blot using LiCOR fluorescence intensity measurements revealed that VAPB levels were unchanged throughout the infection (Fig. 7C).

### Host vesicle fusion proteins accumulate at concentrations of the HSV-1 nuclear egress protein UL34 at and around the NE

Previous studies have shown that HSV-1 nuclear egress is a highly regulated process driven by the heterodimeric viral complex of pUL31 and pUL34 [54, 55]. This complex accumulates at the NE during infection where it both recruits cellular kinases to phosphorylate lamins thus promoting their disassembly so that the virus has access to the inner nuclear membrane for fusion events and appears to also directly participate in primary envelopment [7, 14, 56]. VAPB exhibited almost complete co-localization with pUL34 at the NE (Fig. 8A)

**Figure 8.**
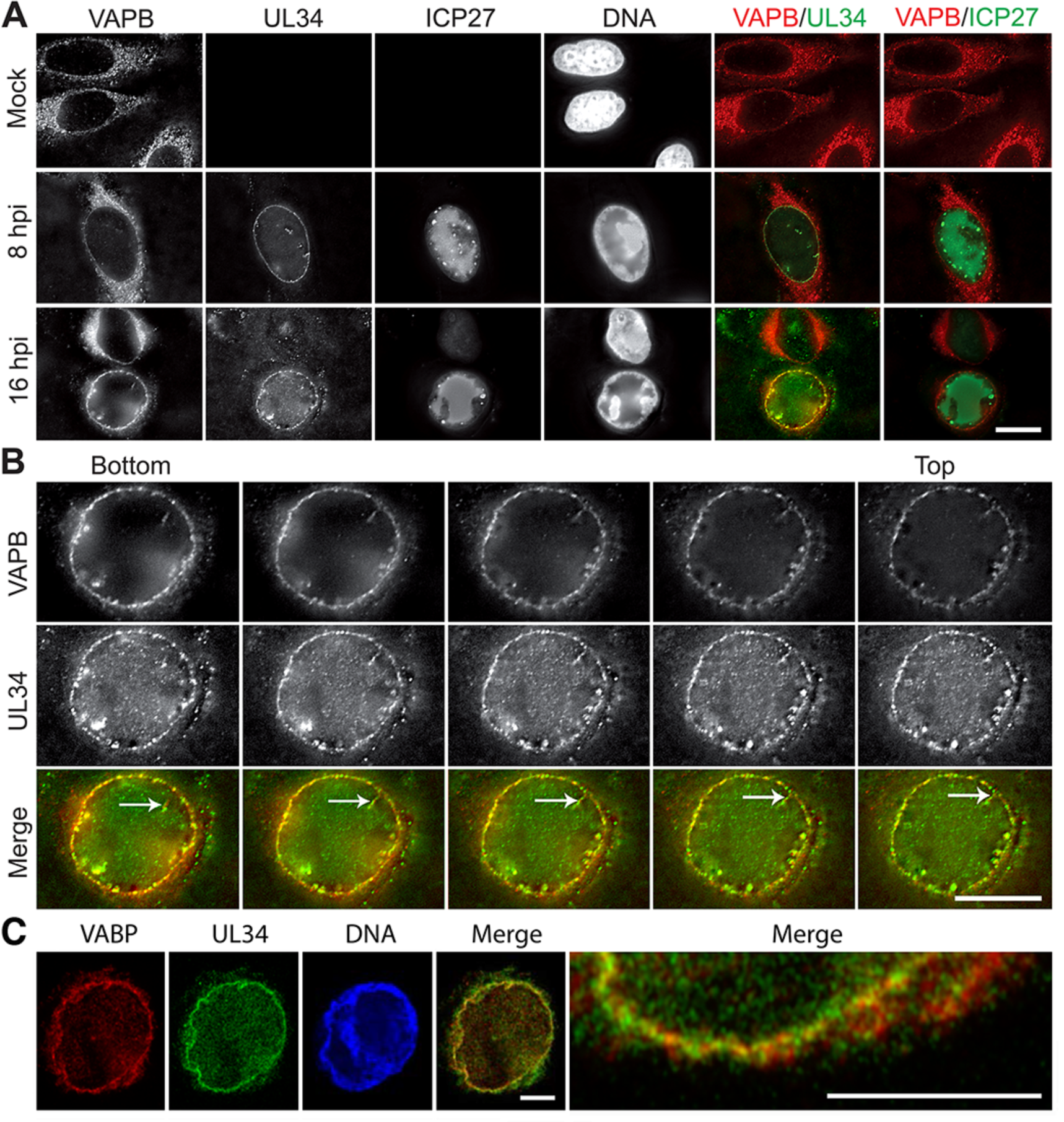
VAPB colocalizes with pUL34 at the nuclear membrane of HSV-1 infected cells. (**A**) HeLa cells infected with HSV-1 vBSGFP27 (encoding WT ICP27 fused to GFP) at MOI 10 were fixed at 8 and 16 hpi, permeabilized, and stained with pUL34 and VAPB antibodies. VAPB, but not control ICP27, co-localized with pUL34. Scale bar, 10 μm. (**B**) As internal punctae were observed in addition to NE staining, z-series were also taken to determine whether VAPB punctae were from nuclear invaginations or other non-membrane associated structures. All images are deconvolved from stacks with individual sections shown. Sections (0.2 μm each step) clearly show that internal punctae (single arrows) are connected in different sections to the membrane. Scale bar, 10 μm. (**C**) HeLa cells infected with wild-type HSV-1 strain 17+ were fixed at 10 hpi and stained with VAPB mouse monoclonal antibodies, pUL34 chicken antibodies and DAPI. Images were taken using a Zeiss 880 confocal Airyscan super resolution microscope. Considerable co-localization was observed between VAPB and pUL34. Scale bars, 5 μm.

The specificity of the co-localization with pUL34 was underscored using an HSV-1 strain in which ICP27 WT protein was tagged with GFP. ICP27 is both involved in nucleocytoplasmic trafficking of viral intronless mRNAs and targets host cell intron-containing RNAs for destruction [30, 57, 58]. Accordingly, ICP27 accumulates partly at the NE and also in nucleoplasmic punctae; however, these are different from the sites of primary envelopment as ICP27 is also known not to be in the capsid or tegument of primary particles. No overlapping fluorescence signals were observed at all between ICP27 and VAPB (Fig. 8A, most right panel).

As a transmembrane protein VAPB should in theory only be in the NE and not in the nucleoplasm. Yet some of the staining signal appeared to be several microns away from the NE, appearing to occur in punta in the nucleoplasm. This type of staining pattern has previously been reported for pUL34 as infection often induces extensive invaginations of the NE [55, 59]. To confirm that VAPB punta are from invaginations 0.2 μm sections were taken in imaging for several cells. Continuity to the membrane was observed when following puncta through individual sections, revealing these internal punctae to be membrane invaginations of the NE as well as showing co-localization between VAPB and pUL34 throughout (Fig. 8B). Co-localization was also observed for both punta and at the membrane using super resolution microscopy (Fig. 8C).

### VAPB accumulates in both inner and outer nuclear membranes

VAPB could be involved in either envelopment or de-envelopment or both. If just involved in de-envelopment it might be expected to only accumulate in the ONM and not translocate to the INM. Thus its distribution between the membranes was investigated in both mock infected and HSV-1 infected cells by immunogold labelling electron microscopy. Although typically only 3-5 particles were observed in the region of the NE captured in a particular section, nearly all images examined had particles in both inner and outer nuclear membranes in both the mock infected (Fig. 9A) and HSV-1 infected (Fig. 9B) cells. The finding of VAPB in the NE of mock-infected cells is consistent with its identification in NEs by the mass spectrometry data (Table 2 and Supplemental Table S1). Gold particles at the NE were counted, considering separately those in the ONM, the INM, the NE lumen, and at the NPC (Fig. 9C). This indicated an increase in the INM pool during HSV-1 infection, consistent with the NE increase observed by immunofluorescence microscopy (Fig. 7A). Most strikingly, however, was the capturing in several images of virus particles inside the nucleus (possibly touching the membrane above the section) that contained gold particles labelling the VAPB (Fig. 9D and E). This strongly argues that VAPB normally participates in the process of primary envelopment.

**Figure 9.**
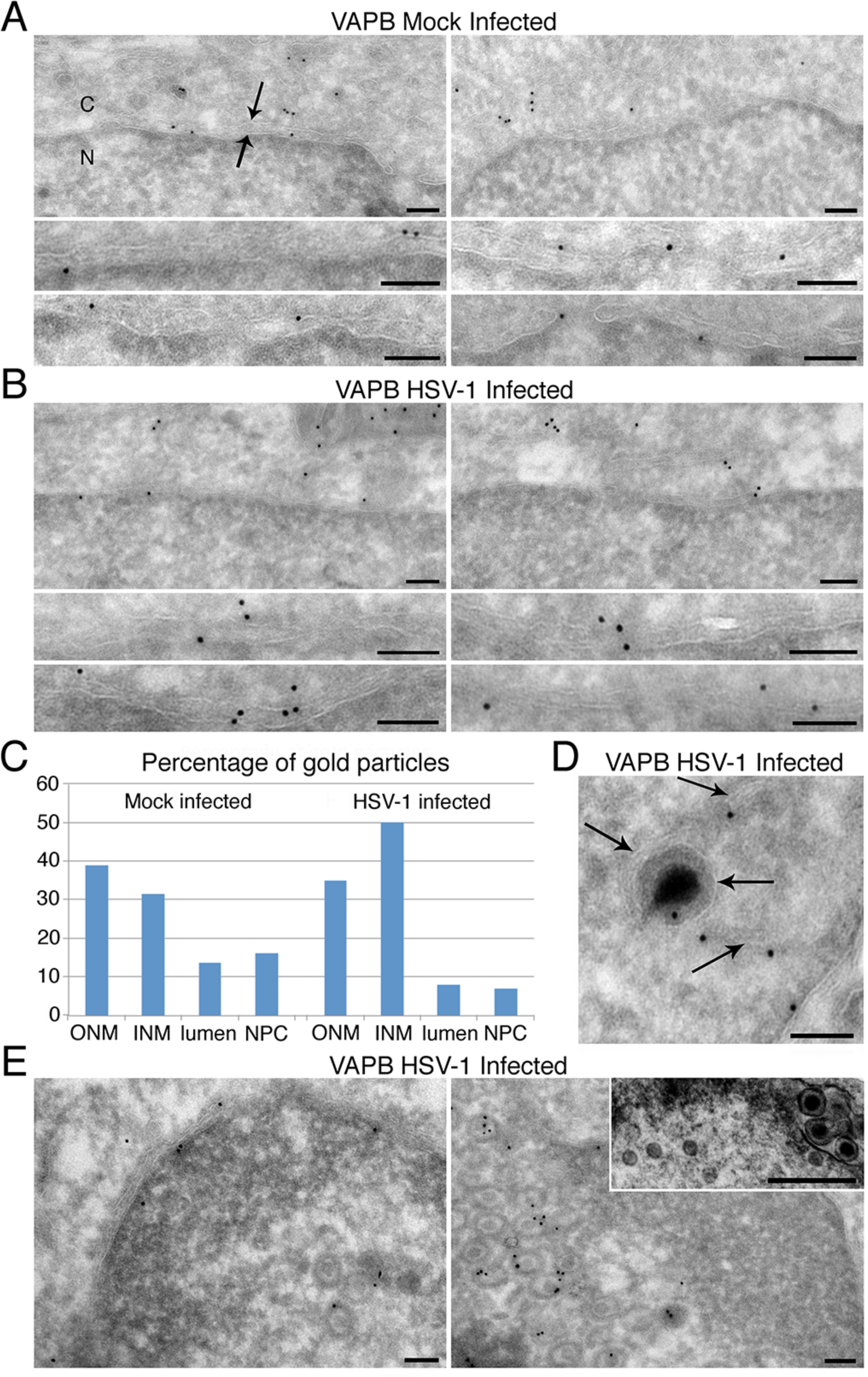
VAPB accumulates in both the ONM and INM. Hela cells either mock infected or infected with WT HSV-1 MOI 10 for 16 h were fixed and cryosectioned prior to labelling with VAPB mouse monoclonal antibodies and anti-mouse conjugated gold particles for electron microscopy. In all panels the cytoplasm is on the top and the nucleoplasm is on the bottom with this labeled in the upper left panel by C and N respectively. This panel also contains two arrows with that facing down delineating the ONM and that facing up delineating the INM. All scale bars, 100 nm. (**A**) Mock infected cells. Note the lower right panel contains one gold particle in transit at the NPC and one in the INM. (**B**) HSV-1 infected cells. (**C**) Quantification of 206 and 163 gold particles at the NE from respectively the mock infected and HSV-1 infected populations. The percentage of total NE particles touching the ONM and the INM are given along with those in the NE lumen and those at the NPC are given. (**D**) High magnification image of a most likely HSV-1 primary enveloped particle with a gold particle indicating VAPB just under the primary envelope membrane. Note that this appears to be located in a membrane bound invagination, possibly from the NE lumen as a membrane can be distinguished within the nucleoplasm (rightward pointing arrows). This membrane also is associated with gold particles. The presumed primary envelope is indicated by the sole leftward pointing arrow. (**E**) Lower magnification images showing multiple nuclear associated virus particles labelling with the VAPB immunogold particles. Although it cannot be fully ascertained from these images if these particles have a primary envelope, comparison with the inset image from standard electron microscopy of the VAPB knockdown infected cells shows that they have more the appearance of the non-enveloped (left) than enveloped (right) particles.

### Discussion

The data reveal for the first time that HSV-1 utilizes a class of host proteins, the vesicle fusion proteins in viral nuclear egress. The cellular processes of vesicle transport and vesicle fusion are extremely complex, involving many proteins for even simplistic modeling and dozens more for functional pathways according to genetic and interaction data [19, 20]. Though herpesviridae are among the larger virus genomes, it would require devoting a significant percentage of their genome to cover the range of functions in these complex events. Thus, despite recent arguments that the pUL31 and pUL34 virus-encoded proteins known to be important for nuclear egress are sufficient for this process [12, 13], it seems likely that host proteins are used at least for more efficient egress. Herpesviruses frequently co-opt host cell machinery to support various aspects of their life cycle [21-23] and accordingly, the involvement of ESCRT pathway proteins has been recently shown for Epstein-Barr virus (EBV) [60]. In this case, the Chmp4b protein important for scission complex assembly largely co-localized in perinuclear aggregates with EBV protein BFRF1 and inhibition of the Alix bridging protein resulted in accumulation of capsid proteins in the nucleus [60]. However, they also reported that the ability of the EBV BFRF1 protein to induce vesicle formation was not found for HSV-1 pUL34, suggesting that HSV-1 requires additional mechanisms for the initial envelopment step [60]. Thus the identification of VAPB, Rab11b and Rab18 here as contributing to virus egress is not surprising. What is perhaps surprising is the extent of co-option of host cell vesicle fusion machinery as other vesicle fusion proteins VAMP7, Rab9a, Rab2A and STX7 all yielded similar NE localization and an increase in the HSV-1 infected MMs compared to mock-infected MMs and thus might also contribute to herpesvirus egress.

VAPB is a member of the Vesicle-associated membrane protein (VAMP)-Associated Protein family of ER C-tail anchored proteins. It functions as an adaptor protein to recruit target proteins to the ER and execute various cellular functions such as lipid transport, membrane trafficking and membrane fusion [61]. A nuclear link has also previously been found for VAPB where its knockdown resulted in a failure of NPC proteins gp210 and Nup214 and the nuclear membrane protein emerin to accumulate at the nuclear envelope [42]. However, as VAPB was predominantly found in the ER in this study, this could as easily reflect general disruption of protein synthesis, post-translational modifications and trafficking caused by VAPB knockdown which is known to induce Golgi fragmentation and disrupted ER to ERGIC (ER-Golgi Intermediate Compartment) to Golgi trafficking [62, 63]. In contrast, we found clear accumulation of VAPB in the nuclear membrane during HSV-1 infection and its considerable co-localization with pUL34, suggesting a direct function in nuclear egress. This does not discount, however, the possibility that VAPB has both direct and indirect effects on HSV-1 processing and trafficking or a possible additional role in secondary envelopment.

A potential additional role for the two Rab proteins tested in secondary envelopment is also a possibility, especially as a previous study reported Rab11b involvement in secondary envelopment [53]. Rab proteins are membrane-associated proteins belonging to the Ras-related small GTPases. There are over 50 Rab proteins encoded by mammalian genomes with only 19 of them having functions in the Golgi and only 6 of these functioning exclusively in the Golgi [64]. Moreover, Rab11b is reported also in the recycling and early endosomes [65, 66] and Rab18 is reported also in the ER, endosome and lipid droplets [67, 68]. Functionally, Rab proteins have been shown to be crucial regulators for many aspects of membrane trafficking including vesicle formation, budding, docking and fusion so that they could contribute to multiple aspects of both envelopment and de-envelopment, as well as secondary envelopment.

The only previously known functional links between Rab11b and viruses are a study showing its involvement in herpes secondary envelopment in the Golgi [53] and a report of its involvement in hantavirus release from cells [69]. Rab18 in contrast is linked to multiple RNA viruses for their assembly around lipid droplets in the cytoplasm, including hepatitis C and Dengue virus [70, 71]; however, such functions are clearly distinct from its contribution to herpesvirus nuclear egress. Nonetheless, due to the wide range of functions linked to Rab proteins it remains possible that some of their effects on the virus could be indirect. For example, Rab11b was recently shown to be involved in proper sorting of the protease-activated receptor-1 protein [72], thus raising the possibility that Rab protein recruitment of as yet unknown proteins to the NE could contribute to post-translational processing of herpesvirus proteins.

One critical question that remains about the role of these vesicle fusion/trafficking proteins is whether they contribute to primary envelopment or de-envelopment or both during nuclear egress. The immunogold labelling electron microscopy data showed VAPB associating with nucleoplasmic virus particles, making a role in primary envelopment almost certain. However, the immunogold electron microscopy data also clearly shows VAPB localizes both to the ONM and the INM during HSV-1 infection, which indicates that it is present to potentially contribute to either or both steps. It is important to note that while the appearance in either membrane could reflect functionality, if these transmembrane proteins translocate by lateral diffusion in the membrane through the peripheral NPC channels similarly to other nuclear envelope transmembrane proteins [24, 25, 27-29], then the presence in both membranes could also simply reflect this diffusion process due to the many NPCs in the membrane. In considering a possible dual function, it is noteworthy that previous studies of gB and gH depletion or mutants revealed an accumulation of primary enveloped particles in both the nucleoplasm and in the lumen of the nuclear envelope [17, 18]. This is a similar phenotype to that of US3 deletions and would be the expected phenotype for a role in the de-envelopment phase of nuclear egress [14, 15]. In contrast, knockdown of VAPB, Rab11b and Rab18 all resulted principally in accumulation of non-enveloped encapsidated virus particles in the nucleoplasm with very few observed in the nuclear envelope lumen. This is more what would be expected for a role in primary envelopment at the inner nuclear membrane and, accordingly, is consistent with the observation of VAPB in both the INM and ONM during normal infection as any protein in the INM that got into primary enveloped particles would be redistributed to the ONM upon de-envelopment. It is important to note that this does not contradict any of the data in recent papers arguing that pUL31 and pUL34 are sufficient for primary envelopment as these studies only showed that membrane invaginations can be induced by mixing these proteins with liposomes in one study [12] and overexpressing them in the context of a cell that still has all these vesicle fusion proteins at lower abundance in the non-infected nucleus [13]. Neither study addressed the issue of efficiency compared to a wild-type infection and, though with greatly reduced titers, HSV-1 has been shown to still get out of the nucleus in the absence of these proteins [14-16, 73, 74].

Our findings also re-open the debate on the composition of primary enveloped particles. Although only gD of the viral coat glycoproteins had been identified in the previous study of primary virions [48], other studies have indicated an association of gB and gC [49, 50] that we found of similar abundance to gD in our mass spectrometry analysis. Thus, it seems likely that our datasets contain most or all of the proteins — both viral‐ and host cell-encoded — that are in primary virions. While we cannot clearly distinguish which proteins were present due to synthesis in the ER and which are in captured primary virions, the grouping of higher abundance glycoproteins that were all previously reported in primary enveloped virions suggests that other virion proteins may be identified by similar abundance groupings. Importantly, the visualization of what appear to be primary enveloped particles by electron microscopy in the HSV-1 infected MM preparation suggests that, though daunting considering that 15 L of culture were required per sample for this study, it may be possible to isolate primary enveloped particles from larger MM preparations.

While capsid and some tegument proteins are expected in the primary virions, the functional significance, if any, for glycoproteins in nuclear egress is unclear. However, if they are required for entry into the cell and/or secondary envelopment they could also play a role in primary envelopment and nuclear egress. A clue may come from the colocalisation of gB, gH, pUL31 and pUL34 at the NE together with cellular proteins CD98hc and β1-intergrin. The cellular components of this complex seem to play a role in HSV nuclear de-envelopment [75]. Such viral protein complexes may control membrane fusion events through recruiting the vesicle fusion proteins needed for both primary and secondary envelopment.

Alternatively, viral glycoproteins may enable less specific interactions with a range of NE transmembrane proteins for as yet undetermined functions or to facilitate proximal access of particles to the membrane and retention for envelopment of tegument-coated virus capsids diffusing through the nucleus [76]. The NE actually contains hundreds of different transmembrane proteins [31, 32, 77] that either need to be targeted to get out of the virus path as has been shown for emerin and LBR [78-80] or might also be co-opted by the virus.

Although only VAPB, Rab11b and Rab18 were directly tested for INM accumulation and effects on HSV-1 egress, additional vesicle fusion proteins, VAMP7, Rab9a, Rab2A and STX7 based on the mass spectrometry data also accumulated in the HSV-1 infected MMs compared to mock-infected MMs. This argues for considerable redundancy in the system that can readily explain why combined knockdowns did not yield a greater reduction in viral titers. However, it will also make more complex the task of deciphering the complex interplay of the vesicle fusion process in herpesvirus egress indicated by this work and the previous work with ESCRT proteins [60]. There are many questions still to be answered. What is their relative contribution to primary vs secondary envelopment? What viral proteins recruit the vesicle fusion proteins and do they directly interact? Unfortunately, the addition of tags for pulldowns strongly inhibits the function of the pUL31 and pUL34 proteins and the same applies for most small GTPases tested, making it likely that tags would similarly disrupt functional interactions of the Rab proteins. Nonetheless, the specific targeting of these vesicle fusion proteins to the INM during infection and the fact that other proteins from different viral stages were unaffected by the vesicle fusion protein knockdowns argues that the function of the vesicle fusion proteins is focused on the specific processes of envelopment and/ or de-envelopment.

## ACKNOWLEDGEMENTS

We thank Richard Roller for UL34 antibody, Oscar Miller for VAPB antibody, and Thomas Mettenleiter and Barbara Klupp for the US3 antibody. We also thank Frazer Rixon for EM support for images of MM preps, Helen Grindley for assistance with the immunogold labelling, and Alexandr Makarov for support in NE and MM preparations. N.S.R. was supported by a University of Edinburgh Principal’s Studentship, C.D was supported by Wellcome Trust studentship 109089, S.V.G. was supported by the Medical Research Council. S.K.S and L.F. are supported by the Stowers Institute for Medical Research. This work was supported by Wellcome Trust Senior Research Fellowship 095209 to E.C.S. and the Wellcome Trust Centre for Cell Biology core grant 092076.

## Supporting Information Legends

Table S1 is an excel file listing all peptides identified in the NE and MM +/- HSV-1 infection mass spectrometry datasets. Two additional sheets are attached to the main Table S1 excel file that contain more details of the protein identifications highlighted in Tables 1 and 2 from the text.

